# Engineering a novel probiotic toolkit in *Escherichia coli Nissle1917* for sensing and mitigating gut inflammatory diseases

**DOI:** 10.1101/2024.06.17.599326

**Authors:** Nathalie Weibel, Martina Curcio, Atilla Schreiber, Gabriel Arriaga, Marine Mausy, Jana Mehdy, Lea Brüllmann, Andreas Meyer, Len Roth, Tamara Flury, Valerie Pecina, Kim Starlinger, Jan Dernič, Kenny Jungfer, Fabian Ackle, Jennifer Earp, Martin Hausmann, Martin Jinek, Gerhard Rogler, Cauã Antunes Westmann

## Abstract

Inflammatory Bowel Disease (IBD) is characterized by chronic intestinal inflammation with no cure and limited treatment options that often have systemic side effects. In this study, we developed a target-specific system to potentially treat IBD by engineering the probiotic bacterium *Escherichia coli Nissle 1917* (EcN). Our modular system comprises three components: a transcription factor-based sensor (NorR) capable of detecting the inflammation biomarker nitric oxide, a type 1 hemolysin secretion system, and a therapeutic cargo consisting of a library of humanized anti-TNFα nanobodies. Despite a reduction in sensitivity, our system demonstrated a concentration-dependent response to nitric oxide, successfully secreting functional nanobodies with binding affinities comparable to the commonly used drug Adalimumab, as confirmed by ELISA and in vitro assays. This newly validated nanobody library expands EcN therapeutic capabilities. The adopted secretion system, also characterized for the first time in EcN, can be further adapted as a platform for screening and purifying proteins of interest. Additionally, we provided a mathematical framework to assess critical parameters in engineering probiotic systems, including the production and diffusion of relevant molecules, bacterial colonization rates, and particle interactions. This integrated approach expands the synthetic biology toolbox for EcN-based therapies, providing novel parts, circuits, and a model for tunable responses at inflammatory hotspots.

**Graphical abstract:** 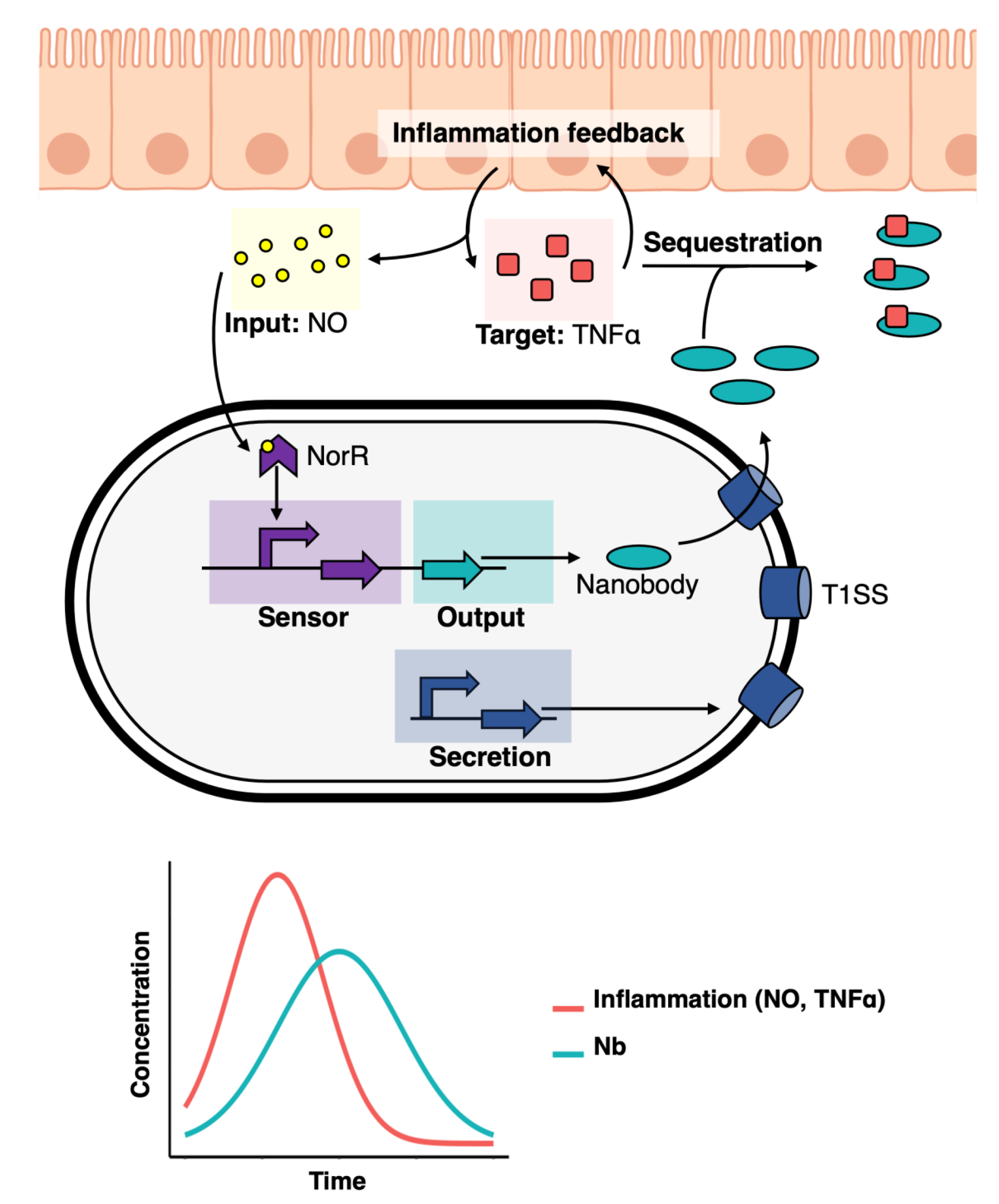

**Graphical Table of Contents**. **The engineered probiotic system**: Inflamed intestinal cells release the inflammatory regulator TNFα (depicted as red squares), which promotes inflammation through a positive feedback loop. Concurrently, these cells produce large amounts of nitric oxide (NO, represented by yellow circles) during inflammation. Our custom-engineered EcN biosensor can detect NO using a NorR-based sensor (in purple) and subsequently trigger the production of nanobodies (in turquoise). These nanobodies are then released into the extracellular environment via a specially engineered secretion system in the bacterial host (shown in dark blue). Once outside the cell, the nanobodies attach to TNFα, effectively sequestering them and reducing inflammation. The graph at the bottom of this panel illustrates the general behavior of our system: nanobody production starts upon reaching a certain NO concentration threshold and continues in an NO-dependent fashion. As nanobodies are produced, they capture TNFα, leading to a reduction in inflammation and a decrease in NO production. This decrease in NO then halts the nanobody production.

**Significance:** Probiotics can be engineered to detect and act upon extracellular disease indicators, optimizing therapeutic outcomes. Particularly, self-regulating sense-and-respond genetic circuits have the potential to enhance the accuracy, efficacy, and adaptability of treatment interventions. In this study, we developed and characterized a new integrated and modular toolkit that detects a gut inflammation biomarker, specifically nitric oxide, and responds to it in an inducible manner by secreting humanized nanobodies targeting the pro-inflammatory molecule TNFα. We also develop a coarse-grained mathematical framework for modelling engineered probiotic activity in the gut. This novel system contributes to current efforts to develop new engineered probiotic systems and holds promise for inspiring new treatments for gut inflammation associated with various autoimmune diseases.

## INTRODUCTION

Inflammatory Bowel Diseases (IBD) are chronic relapsing inflammations of the gastrointestinal tract that affect more than six million people worldwide ^1–5^. Inflammation of the intestinal mucosa compromises barrier function, exposing deeper gastrointestinal layers to luminal antigens and microbiota, which triggers aberrant immune responses and maintains local and systemic inflammation ^2^. Current pharmacological interventions aim to induce clinical remission by reducing mucosal inflammation and alleviating disease symptoms.

Among the approved therapies for IBD^6^, monoclonal antibodies against pro-inflammatory cytokines like tumor necrosis factor (TNFα), IL-12/23, or integrins are particularly effective ^6^. TNFα is a key pro-inflammatory mediator with elevated levels in inflamed gut tissue^7^, making it an attractive drug target with demonstrated therapeutic benefit ^6,8,9^. However, the systemic action of these therapeutics can lead to immunosuppression, increasing the risk of serious infections and lymphoma ^8,10^. Therefore, there is a high demand for new therapeutic solutions that target mucosal inflammation more precisely and are cost-effective ^3,10,11^.

Engineered probiotics^12,13^ offer a potential solution for such treatments, being able to reach inflammatory hotspots in the gut where the mucus barrier is compromised by chronic inflammation ^14^. The probiotic Escherichia coli Nissle 1917 (EcN) ^15–17^ is naturally present in the human gut and has been widely used to treat intestinal diseases due to its anti-inflammatory and antimicrobial properties ^17–24^. Thus, EcN is a promising chassis for targeted gut therapies ^25^. Over the last decade, this strain has been extensively engineered to produce biomolecules at disease sites, particularly for treating intestinal diseases ^26–33^. However, despite recent developments in expanding the tools and biological parts for engineering EcN, there remains a shortage of self-regulating genetic circuits that can recognize specific biomarkers and respond by producing therapeutic molecules^34^.

To address this challenge and contribute to the expansion of the EcN Synthetic Biology toolbox, we designed, engineered, and characterized a new genetic circuit for EcN to act as a biotherapeutic against gut inflammation. This circuit detects nitric oxide (NO) as a biomarker and responds by producing and secreting nanobodies to sequester TNFα and locally reduce inflammation. To date, only one other study has created a similar functional system, however, without a biomarker-induced expression and using alternative components in their circuitry^33^. The scarcity of such systems in EcN highlights the need for alternative systems such as the one presented in our study.

Nitric oxide is a free radical synthesized by inducible nitric oxide synthase in gut epithelial cells, with increased concentrations at inflamed sites ^35,36^. This small molecule can also penetrate bacterial membranes without specialized surface receptors ^37^, making it an effective biomarker for inflammation. In this study, we utilized a NO biosensor endogenous to *E. coli*, specifically the NorR-pNorV system, which was previously modified and characterized by Xiaoyu J. Chen et al.^38^, to trigger the expression of the nanobody delivery system.

Nanobodies, single-domain antibodies that can bind specific antigens^39,40^, are advantageous in therapeutic applications due to their superior tissue penetration, stability, and ease of production by bacteria^33^. These nanobodies can be “humanized” to reduce immunogenicity by modifying specific amino acids ^41^. In this study, we used humanized nanobodies developed by Silence et al. ^42^, producing them for the first time in EcN.

The secretion of nanobodies is essential for TNFα inactivation since this cytokine is present in the gut extracellular environment. Most secretion systems in gram-negative bacteria such as EcN typically release proteins into the periplasmic space rather than the surrounding environment^43^. Thus, we utilized the Type I Hemolysin A Secretion System from uropathogenic *E. coli* ^44–48^. This system has the advantage of being one of the smallest secretion complexes in gram-negative bacteria, and its functionality has not been described in EcN before.

Thus, in this study, we engineered a novel self-regulated system to produce and secrete anti-TNFα nanobodies in response to NO, aiming to reduce intestinal inflammation. Our data shows that although NO sensitivity was lower than reported in a previous study^38^, our system successfully expressed a variety of humanized nanobodies in an inducible manner. We also demonstrate that the produced nanobodies can be effectively secreted to the extracellular environment, retaining their functional capabilities to bind TNFα and reduce inflammation in cell-based assays. This indicates that this system can also facilitate the screening and purification of nanobodies or other proteins of interest in future studies using EcN. Lastly, we developed a mathematical framework to investigate relevant parameters for gut inflammation treatment, addressing the scarcity of modelling tools for such systems.

## RESULTS

### Experimental design

We designed our system by integrating two independent modules on separate plasmids: a sensing module and a secretion module. The sensing module recognizes NO concentrations through the NorR transcription regulator and promotes the production of nanobodies in an inducible manner. The secretion module encodes a secretion system that allows the secretion of nanobodies into the extracellular environment. We characterized each component of our system independently before combining the complete engineered device. This allowed us not only to provide a proof of concept for each subsystem but also to optimize some of them in an iterative process. Firstly, we assessed different architectures of our sensing system through fluorescence reporter-based assays, characterizing their limit of detection and output fold-change in response to different NO concentrations. Secondly, we assessed the production and secretion of nanobodies and their activity using in vitro and cell-based assays. Finally, we tested the whole device and its ability to produce nanobodies in an induced manner. We complemented our study with a simple yet insightful mathematical framework assessing the interactions between the EcN and inflammation sites, focusing on the production rates of NO, the production rates of TNFα and the bacterial response to NO through production of the anti-TNFα nanobodies.

### Characterization of the NO sensing module

To create an inducible system that can sense and respond to inflammation in the gut, we chose a NO-sensitive genetic circuit based on the NorR regulator. NorR is an endogenous transcription factor from *E. coli* responsible for sensing NO concentrations and modulating the expression of genes that are essential for NO detoxification under anaerobic conditions ^49,50^. NorR interacts with NO through a non-haem iron center and binds cooperatively to three enhancer sites at the pNorV promoter to regulate transcription of both *norVW* genes and its own divergently transcribed gene (*norR)* ^49–51^. In *E. coli*, it thereby regulates the activity of the target *norV* gene in a NO-dependent manner. At low NO concentrations, NorR is predominantly present in its free form, which inhibits pNorV. However, at higher concentrations of NO, the radical binds NorR, inducing a conformational change of this protein, which makes it now able to promote σ54-dependent translational activation^52^.

Our sensor was based on a previous study by Xiaoyu J. Chen et al.^38^, consisting of the promoter pNorVβ, an optimized variant of the natural *E. coli K-12* pNorV lacking the second integration host factor (IHF) binding site ^38^. We placed the promoter upstream a bicistronic operon containing genes encoding for a superfolder GFP (*sfGFP*)^53^ and for the NorR regulator (*norR*), in this order. The regulatory logic is based on a positive feedback loop that modulates NorR availability in a NO-dependent manner ^38^ (**Figure 1a**, see **Supplementary Methods** and **Supplementary Figures S1-S3** for more information about constructs and plasmids). This architecture ensures low inhibitory NorR levels in the cells but high availability of activated NorR in environments with a high NO concentration ^38^. Due to the potential cellular toxicity of NO^54^, we verified that the concentrations used did not influence EcN cell growth in our experiments (**Supplementary Figure S4**). Removal of the positive feedback loop decreases the induced expression of downstream genes (see **Supplementary Figure S5**). To characterize and compare our NO-sensing constructs’ limit of detection and dynamic range, we performed time-lapse fluorescence plate reader assays. We performed these experiments using EcN cells.

**Figure 1.**
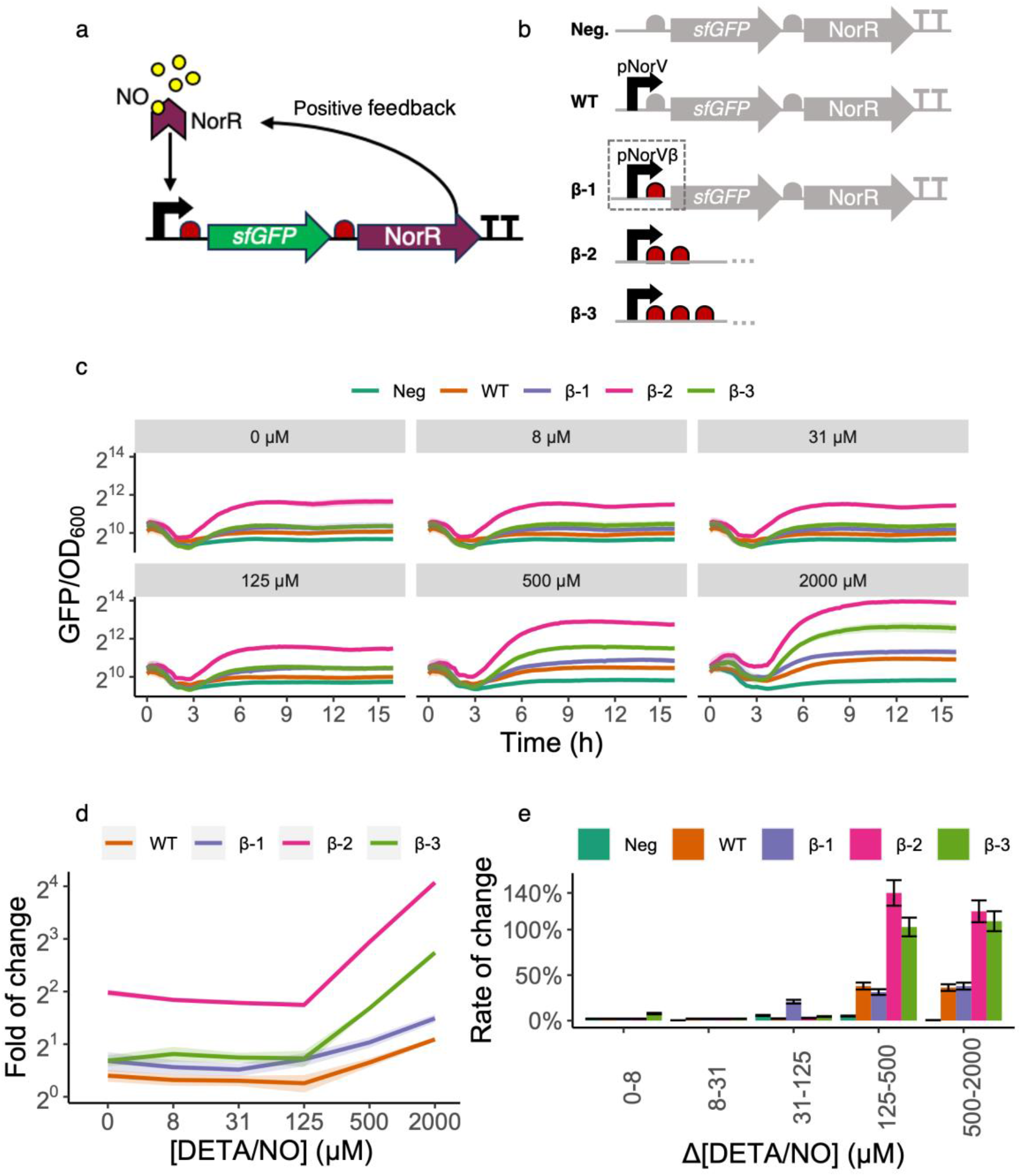
Design and characterization of the NO detection module. **a. NO-dependent activation from NorR.** The NorR transcription factor (represented by the purple chevron) binds its cognate binding site at the promoter pNorVβ (black arrow). When not bound to nitric oxide (yellow circles), NorR acts as a competitive inhibitor of its NO-bound form and represses pNorVβ. However, at high NO concentrations, the NO-bound form of NorR is predominant and acts as a positive inducer of pNorVβ. The presence of *norR* in the inducible operon generates a positive feedback mechanism. Ribosomes are represented in red and the *sfGFP* gene in green. **b. Construct variants characterized.** Our original construct β-1 consisted of *sfGFP* and *norR*, preceded by one RBS each, and placed under the control of the optimized promoter pNorVβ. To avoid read-through, we placed a double-terminator at the end of the operon. We normalized the responses of β-1, β-2, and β-3 to a negative control (Neg) and compared to a positive control (WT). Neg consisted of *sfGFP* and *norR* genes, preceded by one RBS each, and did not contain any promoter, accounting for the intrinsic leakiness of our module. WT consisted of *sfGFP* and *norR* genes, preceded by one RBS each, and placed under the control of the wild-type promoter pNorV.) **c. Time-lapse fluorescence assay for construct characterization.** We have grown each construct for 16 hours (x-axis) on a microplate reader where green fluorescence (arbitrary units) and measured the culture’s OD_600_ every 15 minutes. The y-axis represents normalized fluorescence values (sfGFP/OD_600_). Each panel grid represents a different concentration of diethylenetriamine/nitric oxide (DETA/NO) used to test individual constructs. The DETA/NO gradients we used were 0, 8, 31,125,500, and 2000μM. Each line color represents a construct. Line shadings represent the standard deviation of our biological replicates (*n=3*). We performed all measurements with both biological and technical triplicates. Notice that measurements are on log_2_ scale to facilitate data visualization. **d. Fold of change for each construct.** Each curve represents the fold of change for each construct at T = 8h along a gradient of NO concentrations. Line shadings represent the standard deviation of our biological replicates (*n=3*). Notice that measurements are on the log_2_ scale to facilitate data visualization. **e. Rate of change for each construct.** The bar plots represent the rate of change for each construct for each DETA/NO change of concentration at T = 8h. We calculated rates of change as the relative increase in fluorescence (reported as percentages, y-axis) from an initial NO concentration to the next incremental one. We have performed such calculations for each consecutive pair of concentrations (x-axis). Error bars represent the standard deviation of our biological replicates (*n=3*).

We observed that the NorR circuit design with the best performance in the original study^38^ featured three consecutive ribosome binding sites (RBSs) upstream of the *sfGFP* gene. To investigate the impact of altering the number of consecutive RBSs on the sensitivity of our system, we designed, constructed, and characterized three variants with one, two, or three consecutive RBSs, respectively named β-1, β-2, and β-3 (**Figure 1b**). This approach allowed us to assess the effect of varying the number of RBSs on the sensitivity of our system. To account for background fluorescence, we systematically compared our constructs to a negative control plasmid that did not contain any promoter (**Figure 1b**). We also compared our constructs to the wild-type pNorV with a single RBS (**Figure 1b**).

### The number of Ribosome Binding Sites (RBSs) upstream of *sfGFP* influences its expression levels and the leakiness of the construct

Our first observation was that the pNorVβ system exhibited higher fluorescence levels than the wild-type, regardless of the NO concentration (**Figure 1c**), indicating this system is leakier than the wild-type. By changing the number of RBSs, we observed differences in our detection limits and the overall fluorescent reporter expression. We can observe in **Figure 1c** that our constructs can be increasingly ranked regarding basal sfGFP expression as WT < β-1 < β-3< β-2. Interestingly, the consecutive addition of ribosome binding sites (RBSs) does not result in a linear increase in sfGFP expression. We speculate that this phenomenon may be due to structural consequences arising from repeating sequences in tandem, such as the potential formation of secondary structures or hairpins ^55^. Additionally, ribosome stalling could occur, where ribosomes pause or slow down due to interactions between ribosomes initiated at different RBSs^56,57^.

We observed that β-2 is highly leaky, showing higher sfGFP expression even in the absence of induction ([NO] = 0). The higher expression baseline of β-2 sfGFP expression can also be highlighted in **Figure 1d**, showing the fold-of change in sfGFP expression for each construct at all tested NO concentrations. We also observed in **Figure 1d** that β-1 responds to a lower concentration than the other constructs ([NO] = 125µM). This is further illustrated in **Figure 1e**, which shows the rate of change, a sensitivity metric for each genetic construct to variations in NO levels, as measured by changes in sfGFP fluorescence. The percentage change in sfGFP fluorescence intensity is calculated when the NO concentration shifts from an initial baseline to a new value. This percentage is then normalized against the initial NO concentration, providing a relative measure of change.

### Purified monovalent and bivalent anti-TNFα nanobodies efficiently capture TNFα, comparable to monoclonal antibodies used in the clinics

To develop the nanobody production module, we have selected three previously described anti-TNFα humanized nanobody candidates^42^ and combined these to additionally produce bivalent nanobodies, linked via a short peptide linker (EPKTPKPQPAAA; for monovalent and bivalent nanobodies **see Materials and Methods Table 3**). Firstly, to assess the proper expression and activity of our candidates, we cloned their sequences into the pSBinit^58^ expression vector (**see Materials and Methods, Table 2 and Supplementary Figures S6-S7**), allowing controlled expression upon L-arabinose induction (**see Figure 2a**). We transformed the plasmids into the expression strain *E. coli MC1061* (**see Materials and Methods, Table 1**). After induction, we performed periplasmic extraction for monovalent nanobodies and whole-cell lysis for bivalent constructs (**Figure 2a**) and observed a quantitatively higher output of monovalent nanobodies compared to the bivalent constructs (**Supplementary Figures S8-S9**).

**Figure 2.**
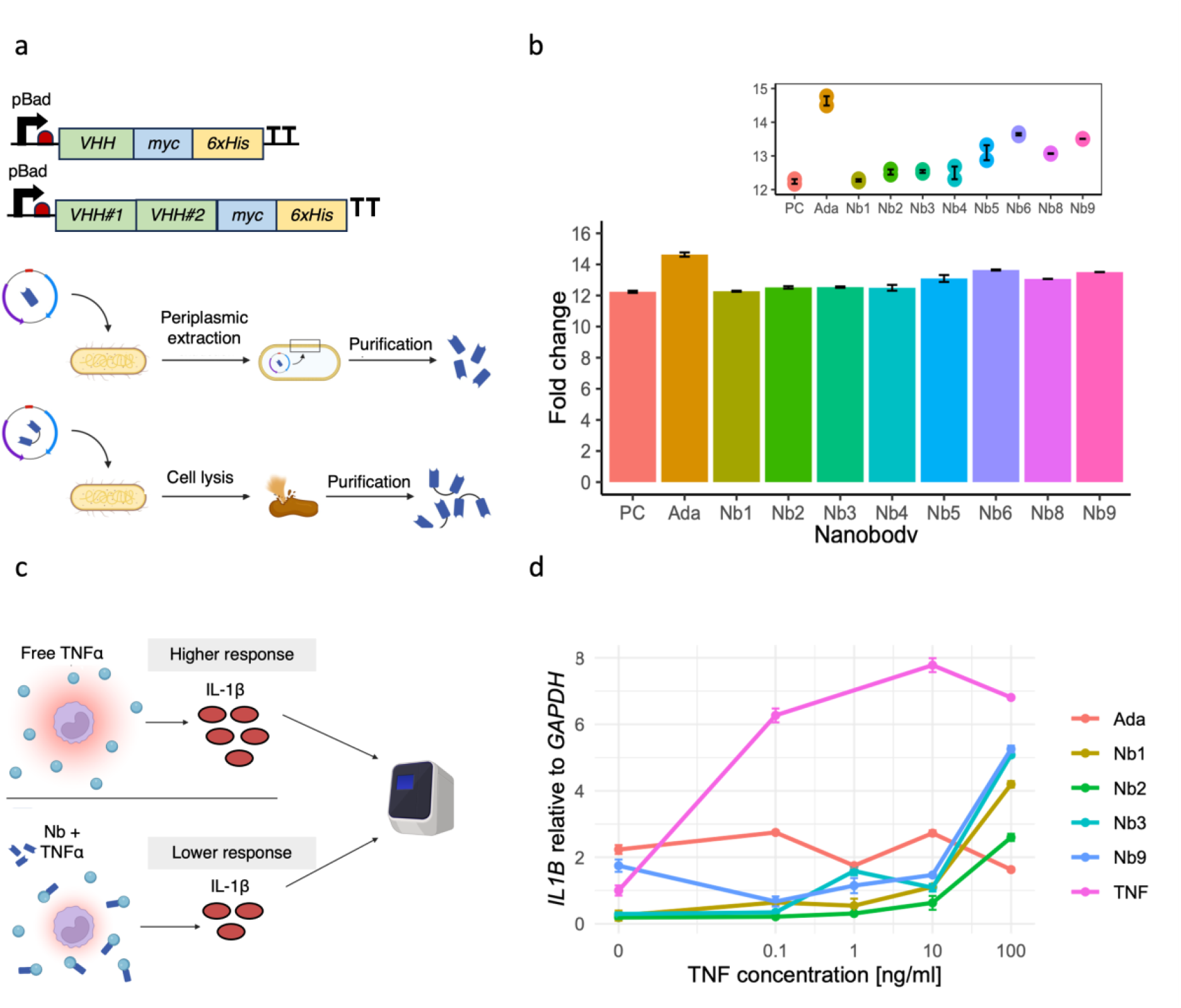
Design and characterization of the purified anti-TNFα nanobodies. a. Design of monovalent and bivalent anti-TNFα nanobodies. We linked bivalent nanobody constructs via a short peptide linker (EPKTPKPQPAAA). To characterize the nanobodies, we added a myc-tag and a his-tag to their C-terminal sites. Their expression was under the control of the inducible pBad system, which relies on the addition of L-arabinose. We induced the expression of nanobodies with the pBad inducible system. We harvested monovalent nanobodies via periplasmic extraction and bivalent nanobodies through whole-cell lysis. We purified all nanobodies by immobilized metal anion chromatography (IMAC). **b. Testing binding capability of purified nanobodies with enzyme-linked immunosorbent assay.** We tested TNFα-binding using an ELISA by capturing the purified nanobodies via their myc-tag. Then, we visualized the binding of nanobodies to biotinylated TNFα with the streptavidin-peroxidase. We measured the absorbance of each well with a plate reader and analyzed the fold change with R studio. **c. Principle of the cell assay used to determine anti-inflammatory properties of purified anti-TNFα nanobodies.** We incubated Human THP-1 monocytes with rTNFα and different purified anti-TNFα nanobodies. We assessed the immune response of the monocytic cell line to rTNFα by quantitatively determining the *IL1B* expression levels with the use of RT-qPCR. The binding of the nanobodies to rTNFα is supposed to inhibit the inflammatory effect observed in untreated but stimulated THP-1 cells. **d. *IL1B* expression compared to *GAPDH* in human THP-1 monocytic cell line.** Quantitative analysis of the inflammatory *IL1B* expression levels revealed a decreased immune response of rTNFα-stimulated cells when purified nanobodies were added, compared to untreated cells (labeled as “TNF”, pink line). Adalimumab is an anti-TNFα monoclonal antibody frequently used in the clinic to treat IBD patients and served in this experiment as a positive control.

**Table 1.**
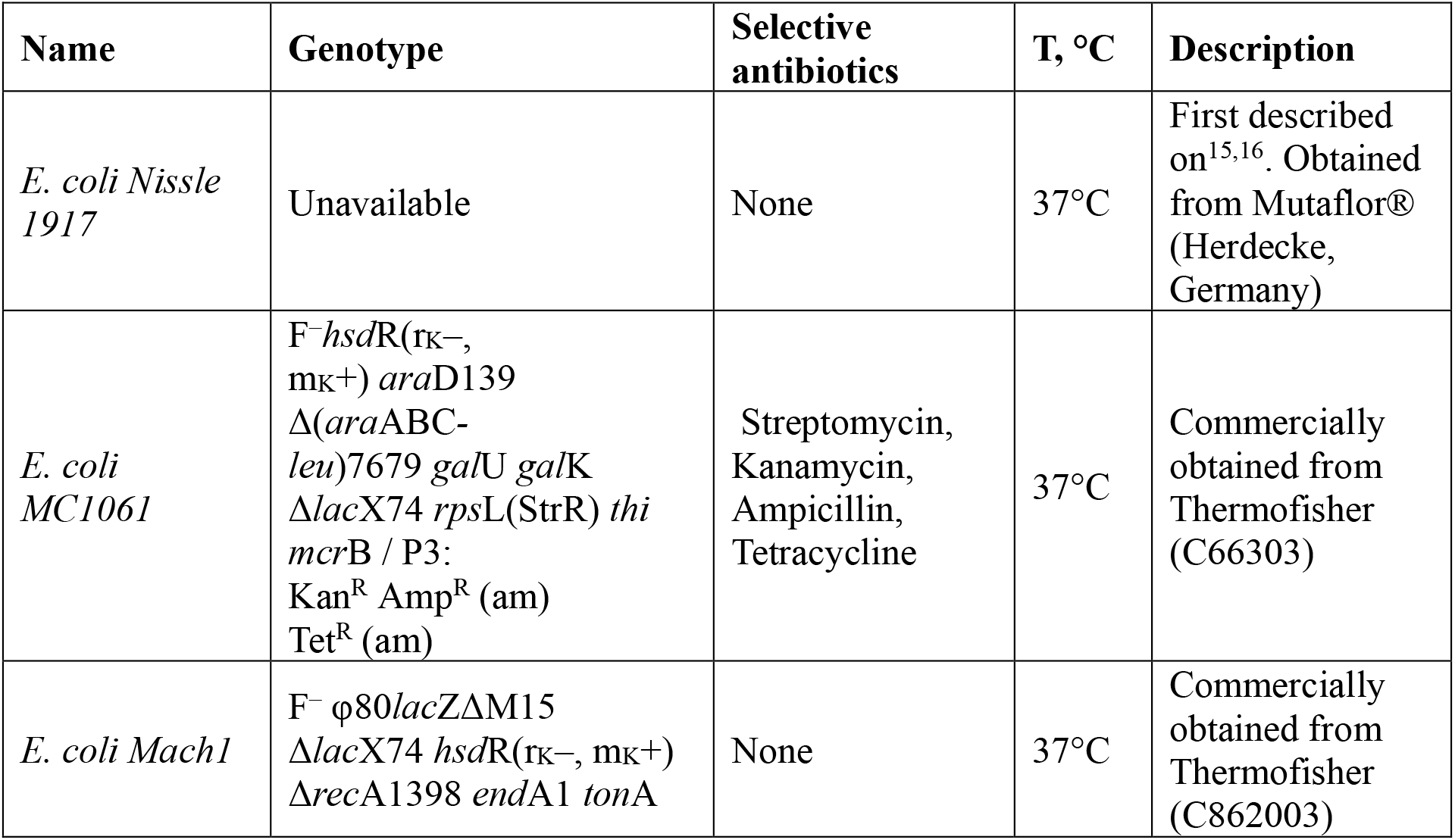
List of bacterial strains used in this study.

**Table 2.** List of plasmids used in this study.

| Name | Description |
| --- | --- |
| piGEM1 | This study. Negative control. Encodes for *sfGFP* and *norR*, does not contain any promoter. |
| piGEM3 | This study. Encodes for *sfGFP* and *norR* under the control of the already characterized inducible promoter pNorV. |
| piGEM2.1 | This study. Encodes for *sfGFP* and *norR* under the control of the inducible promoter pNorVβ. This construct contains 1 RBS directly upstream of *sfGFP*. |
| piGEM2.2 | This study. Encodes for *sfGFP* and *norR* under the control of the inducible promoter pNorVβ. This construct contains 2 RBS and a spacer upstream of *sfGFP*. |
| piGEM2.3 | This study. Encodes for *sfGFP* and *norR* under the control of the inducible promoter pNorVβ. This construct contains 3 RBS and a spacer upstream of *sfGFP*. |
| piGEM2.2N | This study. Encodes for *sfGFP* under the control of the inducible promoter pNorVβ. This construct contains 2 RBS and a spacer upstream of *sfGFP*. This construct does not contain *norR*. |
| pSBinit | Retrieved from <sup>58</sup>, Addgene #110100. E.coli entry and expression vector for FX cloning system, N-terminal pelB signal sequence and C-terminal myc and 6xHisTag |
| purNb1 | This study. Nanobody candidate VHH#2B cloned into pSBinit expression vector via FX cloning |
| purNb2 | This study. Nanobody candidate VHH#3E cloned into pSBinit expression vector via FX cloning |
| purNb3 | This study. Nanobody candidate VHH#12B cloned into pSBinit expression vector via FX cloning |
| purNb4 | This study. Nanobody candidate VHH#2B-VHH#2B cloned into pSBinit expression vector via FX cloning |
| purNb5 | This study. Nanobody candidate VHH#3E-VHH#3E cloned into pSBinit expression vector via FX cloning |
| purNb6 | This study. Nanobody candidate VHH#12B-VHH#12B cloned into pSBinit expression vector via FX cloning |
| purNb7 | This study. Nanobody candidate VHH#2B-VHH#3E cloned into pSBinit expression vector via FX cloning |
| purNb8 | This study. Nanobody candidate VHH#2B-VHH#12B cloned into pSBinit expression vector via FX cloning |
| purNb9 | This study. Nanobody candidate VHH#3E-VHH#12B cloned into pSBinit expression vector via FX cloning |
| pSS | This study. Plasmid encoding HlyB and HlyD required for the HlyA secretion system under a constitutive promoter (J23100). Contains chloramphenicol resistance gene. |
| pNb | This study. Plasmid encoding HlyA and myc-tag under inducible pBad promoter with restriction sites allowing the cloning of the different nanobodies in front of the two tags. Contains ampicillin resistance gene. |
| pNb1 | This study. Nanobody candidate VHH#2B cloned into pNb plasmid via FX cloning |
| pNb2 | This study. Nanobody candidate VHH#3E cloned into pNb plasmid via FX cloning |
| pNb3 | This study. Nanobody candidate VHH#12B cloned into pNb plasmid via FX cloning |
| pNb5 | This study. Nanobody candidate VHH#3E-VHH#3E cloned into pNb plasmid via FX cloning |
| pNb7 | This study. Nanobody candidate VHH#2B-VHH#3E cloned into pNb plasmid via FX cloning |
| pNb8 | This study. Nanobody candidate VHH#2B-VHH#12B cloned into pSBinit expression vector via FX cloning |
| pNO1_Nb1 | This study. Nanobody candidate VHH#2B cloned into piGEM2.1 plasmid via Gibson |
| pNO3_Nb1 | This study. Nanobody candidate VHH#2B cloned into piGEM2.3 plasmid via Gibson |

**Table 3:**
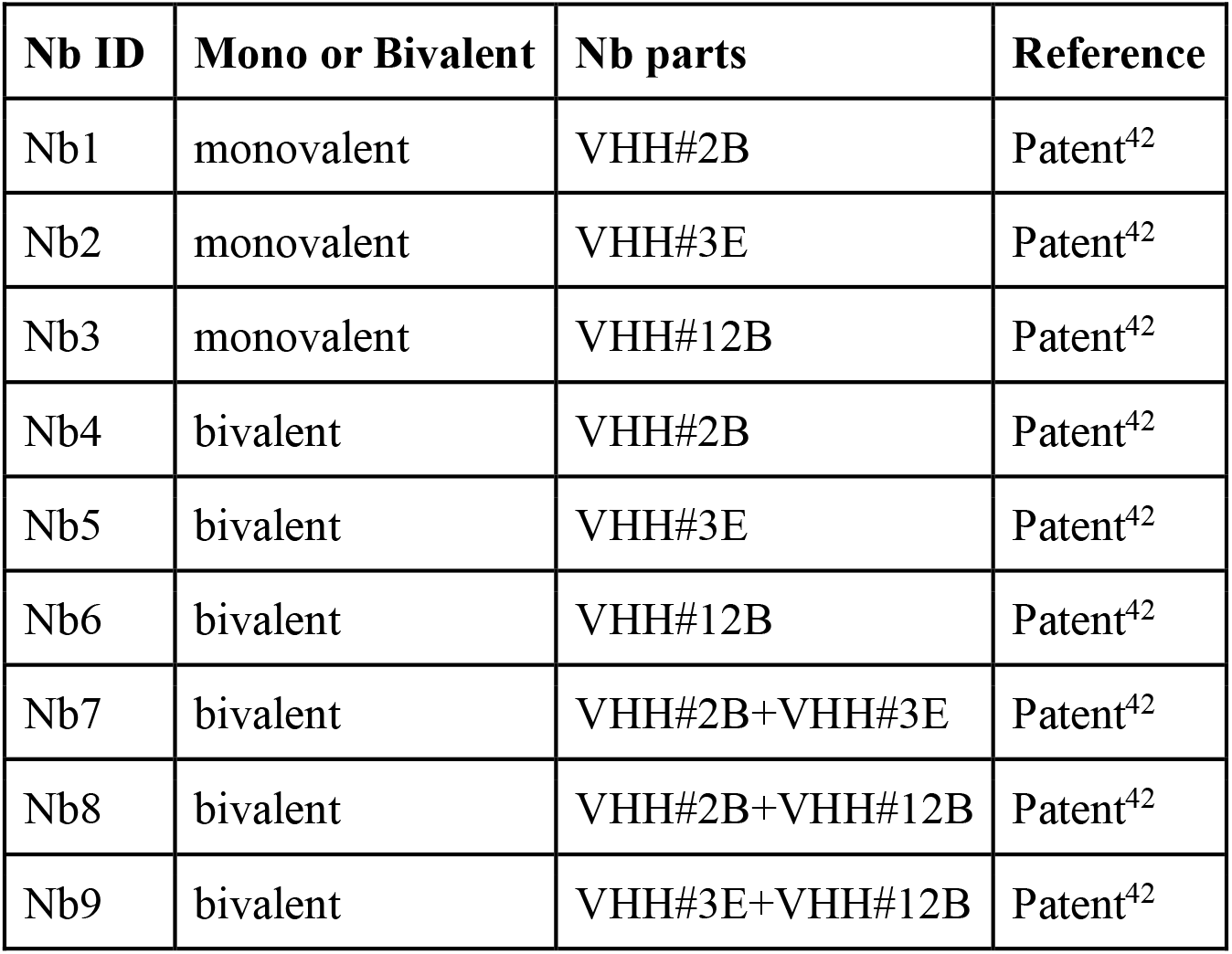
Nanobodies used in this study.

We proceeded by performing an ELISA with the purified nanobodies to test their capability to bind TNFα (**see Materials and Methods**). The **Figure 2b** shows the fold change in the binding capacity of our different nanobody candidates compared to our negative control. The bivalent nanobodies exhibit a statistically significant enhancement in binding efficiency, demonstrating an average 1.3-fold increase over the monovalent nanobodies (see **Supplementary Figure S10**). The bivalent constructs show a mean TNFα binding capacity of 13.5 ± 0.1 (mean ± s.d.), compared to 12.3 ± 0.2 for the monovalent constructs. Adalimumab, an approved monoclonal anti-TNFα antibody that is already used in the clinic to treat IBD (see **Materials and Methods** for antibody purification), is used as a positive control and our bivalent nanobody constructs show a similar binding capability to this therapeutic.

### Anti-TNFα nanobodies show anti-inflammatory effects on stimulated human monocytes in vitro

To evaluate the effect of anti-TNFα nanobodies on the immune response of cells to an inflammatory stimulus in vitro, we stimulated THP-1 human monocytes with increasing concentrations of recombinant TNFα (rTNFα) and subsequently added our purified nanobody candidates (**see Materials and Methods**). We performed real-time quantitative PCR analysis and measured the relative amount of *IL1B* expressed by immune cells as a response to inflammation through TNFα signaling (**Figure 2c**, **Supplementary Methods**). The cytokine IL-1β is an important inflammation mediator and, therefore, a good marker to prove functional TNFα-inhibition^59^.

We were able to observe an up to 4-fold decrease in *IL1B* expression of stimulated monocytes when different nanobodies were added compared to the control cells that only received the inflammatory stimulus (**Figure 2d**). This experiment shows that tested nanobodies have the same capability to lower inflammation as monoclonal antibodies, which are already used in the clinic to treat IBD patients. However, with increasing TNFα concentrations, the anti-inflammatory effect that the nanobodies have on the monocytes seems to slowly decline, indicating that higher concentrations of nanobodies are required to maintain low *IL1B* expression levels. This decline is not observable with the available drug Adalimumab^60^. It is also important to note that the difference between monovalent and bivalent nanobody constructs does not seem to be of great influence on the inflammatory response of triggered monocytes.

### Anti-TNFα nanobodies can be secreted from *EcN*

In order for EcN to deliver nanobodies to its environment, it must be able to secrete them without impacting their function. To achieve this, we engineered the HlyA secretion system^47,48^ into EcN along with fusing the nanobodies to the HlyA-tag, marking them for selective export (**Figure 3a, Supplementary Figure S11**). As a first step, we tested the functionality of the nanobodies after expression and secretion in *E. coli MC1061*. We performed a double transformation of *E. coli MC1061* with two plasmids: our secretion plasmid (**Supplementary Figure S11**) and the pSBinit expression plasmid, which allows for nanobody expression upon L-arabinose induction (**Supplementary Figure S7**). After overnight induction, we harvested the supernatant from the cell culture, and performed a western blot and ELISA to quantify the presence and TNFα binding of the secreted nanobodies (**Figure 3b**). In EcN, the nanobody Nb1 and the bivalent nanobody Nb8 were successfully secreted, and their binding affinities were maintained (**Figure 3c**, (**Supplementary Figures S13-S14**).

**Figure 3.**
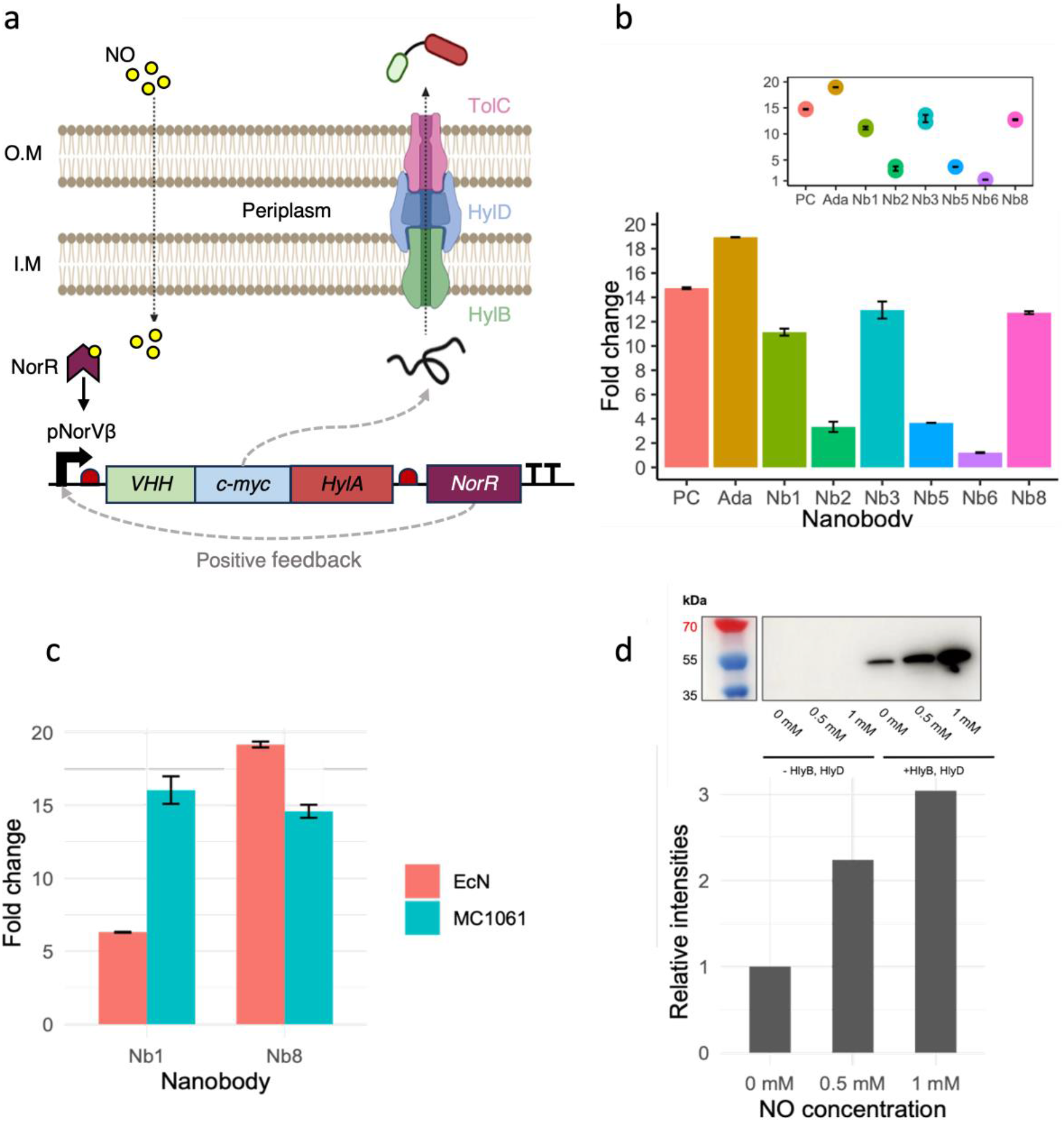
Design and characterisation of arabinose- and NO-induced anti-TNFα nanobodies secretion in *E. coli Nissle 1917* and *E. coli MC1061*. **a. Principle of NO-induced nanobody secretion with the Hemolysin A secretion system.** Nitric oxide is a small organic molecule able to surpass the double membrane of *E. coli*. NO binding to the PnorV-ß promoter induces the expression of the monovalent nanobody candidate Nb1, which is tagged with a myc- and HlyA-tag. NorR expressions result in a positive feedback loop, enhancing the nanobody expression further. Thanks to the HlyA-tag, the produced nanobodies are secreted by the hemolysin A secretion system in a one-step manner into the extracellular space. **b. Arabinose-induced secretion of monovalent and bivalent nanobodies with *E. coli MC1061*.** Western blot and ELISA analysis revealed successful secretion of functional monovalent and bivalent nanobodies upon overnight arabinose induction in *E. coli MC1061*. **c. Arabinose-induced secretion of monovalent and bivalent anti-TNFα nanobodies in *EcN* and *MC1061*.** ELISA analysis shows a successful secretion of functional monovalent Nb1 and bivalent Nb8 nanobodies upon overnight arabinose induction, retaining their TNFα-binding capabilities regardless of the HlyA-tag. **d. NO-induced secretion of monovalent anti-TNFα nanobodies with a single-RBS system in *E. coli MC1061*.** The NO-induced monovalent nanobody secretion was achieved using the single-RBS system (ß-1). This yielded a more dynamic response to NO than the previous two-RBS system (ß-2) (**Supplementary Figure S16**) and a lower baseline expression of monovalent nanobody candidate Nb1 in *E. coli MC1061*. The absence of the two secretion system components (HlyB and HlyD) resulted, as expected, in no secretion of nanobodies. With increasing NO-levels, higher nanobody expression can be observed. A baseline expression in the absence of NO is still present yet weaker than in the ß-2 system (**Supplementary Figure S16**).

### Nitric oxide can be used to trigger anti-TNFα nanobody expression

To create a system capable of sensing NO and thereby triggering the production and secretion of nanobodies, we built a new plasmid, where the monovalent nanobody Nb1 was cloned downstream of the aforementioned pNorVβ promoter (see **Table 2**, **Supplementary Methods and Supplementary Figure S15**). We used the circuit with two RBS (ß-2) upstream of the cloned nanobodies, as it presented the highest expression levels. We used DETA/NO for induction and allowed cells to express nanobodies overnight. We then quantified the presence of nanobodies in the supernatant by western blot, in which we detected secreted nanobodies using a C-terminal myc-tag. The western blot showed that while EcN could sense NO and increase the expression and secretion of anti-TNFα nanobodies, there was still a high-level baseline expression without NO (**see Supplementary Figure S16**).

Despite the high levels of baseline expression, the secreted nanobodies maintained their functionality, as shown in ELISA assays (**see Supplementary Figure S16**). To reduce baseline expression, we tested an alternative circuit differing by having a single RBS (ß-1) upstream of the nanobody coding region. This had previously shown less expression leakage. The single RBS system yielded a more dynamic response to NO concentration in *E. coli* MC1061 after 8 hours of expression. Relative expression showed a threefold increase from baseline to a 1 mM NO concentration (**Figure 3d**). It is worth noting that we observed a basal production of nanobodies even without the addition of the NO inducer (**Figure 3d, rightmost western blot and its corresponding bar plot**). Lastly, the control with no secretion system shows no presence of nanobodies in the supernatant. This confirms the need for a secretion system to export the nanobodies, as cell death does not appear to result in the release of functional nanobodies.

### A coarse-grained model for engineered probiotics in the gut

To support the experimental claims, we constructed a two-dimensional lattice-based reaction-diffusion model ^61–65^ of the gut environment, as in-vivo testing in the gut microbiome is outside the scope of this study. The model is illustrated in **Figure 5a**.

**Figure 4.**
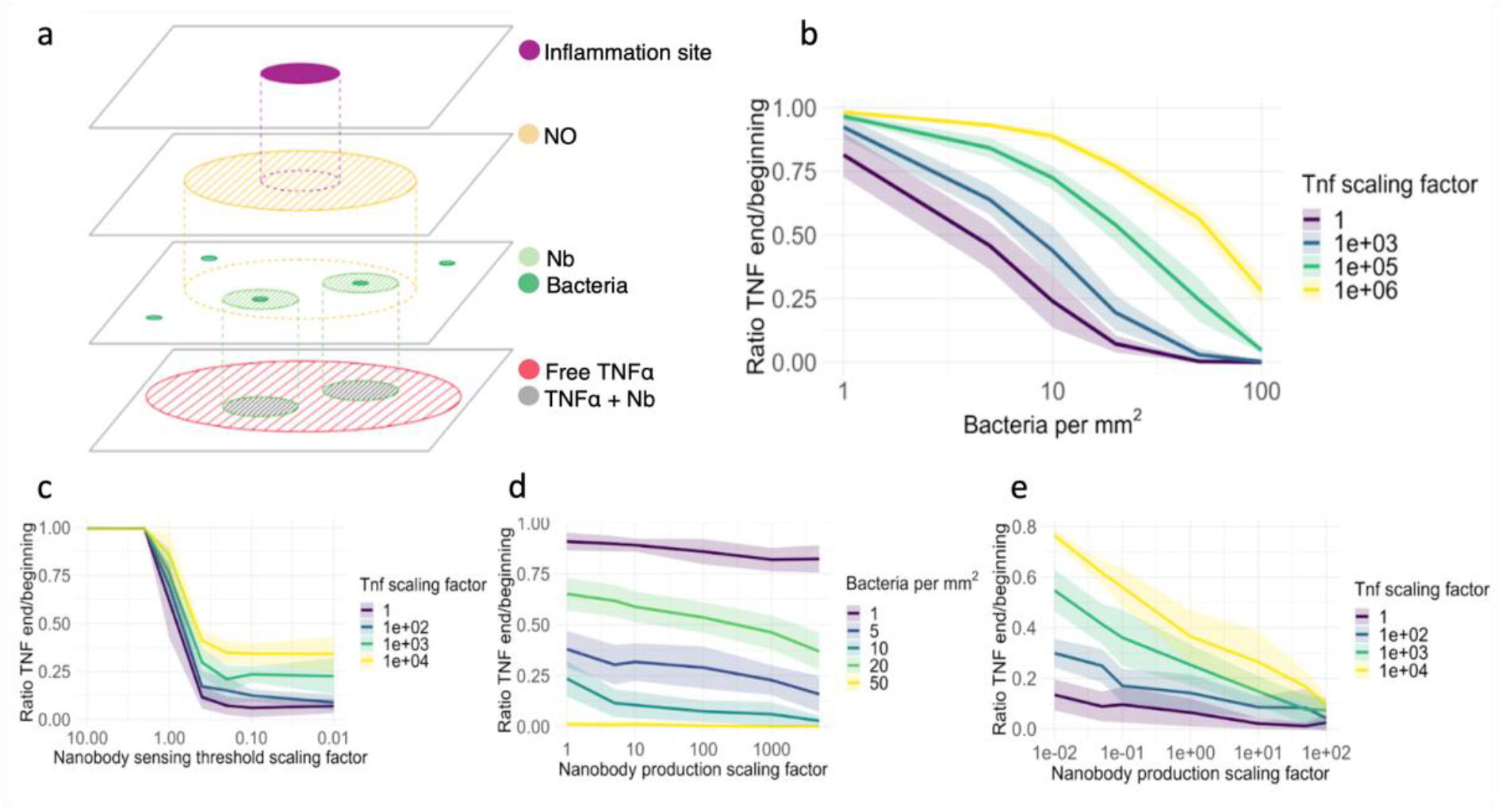
The reaction-diffusion model was evaluated on key parameters. The model’s purpose was to explore which parameters could be essential for the efficacy of our system. Parameters that are not varied are set to their default value, except for the sensing threshold, which we decreased by a factor of 10 during simulations as done in a recent study^38^, for visibility reasons. We simulated each parameter configuration 10 times. Line shadings represent the standard deviation. **a. Illustration of the components of the reaction-diffusion model. b. Relationship between bacterial density and TNFα concentrations. (n = 240). c. Relationship between sensing threshold and TNFα concentrations. (n=280) d. Relationship between nanobody production and bacterial density. (n = 300) e. Relationship between nanobody production and TNFα concentrations. (n = 380)**

The model’s primary objective was to examine the interactions between EcN and inflammation sites in a simplified manner, specifically focusing on the NO concentrations^66^, the production rates of TNFα^67^ and the bacterial response to NO through the production of the TNFα-binding nanobodies ^68^. Model methods and an in-depth description of the parameters used can be found in the model section of the appendix. Through cycles of diffusion, decay, and reemission, we provided a preliminary outlook on the efficacy of our proposed treatment and its potential for healthcare applications. We note that the model is a coarse-grained one, and faithfully representing the gut environment was out of its scope. We focused on identifying the crucial parameters to tune in future work to optimize the treatment before heading into a further testing stage.

To favor interpretability, generalization, and to promote ease of access and collaboration, the model follows a simplicity-based design. The model interprets a 1mm^2^ area of the gut surface as a 2D grid, with each grid cell representing a 1µm^3^ volume, containing the local concentration values for each parameter. The model operates in discrete time steps, with adaptations to make it approach a continuous time scale.

### Estimating the minimum number of bacteria for effective treatment

To get an overview of the importance of the different parameters, we performed a series of simulations where we swept two variables at the same time over the range of our expected values and simulated for 60-second time steps.

In our first series of simulations, illustrated in **Figure 5b**, we estimated the minimum number of bacteria needed to provide effective treatment and evaluated the densities on a great range of biologically plausible TNFα concentrations. The results show that around 20 bacteria per mm² should be enough to cover the inflamed gut area and sufficiently combat the inflammation for the expected TNFα concentrations. For higher concentrations of TNFα, however, the nanobodies produced are not sufficient to combat inflammation, and larger bacterial populations are needed. The graphs suggest a rough relationship of a doubling in bacterial density being able to combat a magnitude higher TNFα concentration. Bacterial density estimates place this requirement at a feasible replacement value of 20 out of 10^4^ gut bacteria per mm² ^69,70^ and thus the required colonization should be feasible and in further experiments we assume that this threshold will be reached. In **Figure 5e**, we investigate whether increasing the nanobody production could also be a viable solution.

### Nitric oxide detection threshold and nanobody production

In our study, the threshold for NO detection, the minimum amount of NO required for nanobody production, is essential for ensuring an inflammation-dependent response. In **Figure 5c**, we evaluated a range of these sensing thresholds and the amount of TNFα at the inflammation sites. As our threshold closely aligns with the expected NO concentration of around 15 μM^36^ (**for details, see Supplementary Methods and Supplementary Figure S17**), even slight decreases in sensitivity lead from the absence of inflammation reduction to complete reduction, even with higher than expected amounts of TNFα. Increased sensitivity of the bacteria towards NO gives diminishing returns, as this mostly affects bacteria in edge regions that detect trace amounts of NO but do not produce nanobodies at the affected location. In turn, there will be an excess production of nanobodies in these regions that hardly contribute to combating inflammation. It is important to note that in our simulations, we kept the amount of NO produced at inflammation sites constant, even for higher TNFα concentrations. In a patient setting, however, NO levels might vary significantly.

### Comparison between bacterial number and nanobody production

In **Figure 5d**, we assessed the importance of the number of bacteria we can introduce against the nanobody production of a single bacteria. For our expected TNFα values, the number of bacteria has a far greater effect than the amount of nanobodies produced per bacteria. This is most likely due to the nanobodies being spread locally, and even at the same amount of net nanobodies produced, greater coverage of gut-inflamed areas ensures that the nanobodies are produced where they need to be. This trade-off also guarantees that no excess amount of nanobodies is produced which could lead to possible side effects. In vivo testing is required to assess the actual viability of our engineered bacteria, and further optimization should be based on this.

In **Figure 5e**, we investigate whether higher TNFα concentrations can be mitigated by increasing nanobody production. The results demonstrate that, with a bacterial density of at least 20 per mm², an increase in nanobody production effectively reduces TNFα levels. However, this effect is less pronounced compared to increasing the number of bacteria, as shown in **Figure 5b**, which more effectively reduces even higher concentrations of TNFα. Nevertheless, increasing nanobody production might be easier to achieve and still offers a viable approach to combating elevated TNFα concentrations.

## DISCUSSION

Here, we describe the development of an integrated molecular system in EcN for the local sensing of gut inflammation and the production/delivery of high-specificity effectors to mitigate such inflammation. Specifically, we have engineered both laboratory and non-pathogenic/probiotic human *E. coli* strains with a coupled system that can secrete nanobodies in a regulated manner upon NO induction. Secretion is achieved through the adoption of an exogenous Type I Hemolysin A Secretion System, which has been characterized in EcN for the first time in this study. We have also characterized a new library of humanized nanobodies in EcN, demonstrating that they can be successfully secreted and retain their functionality in vitro and in cell assays, binding to TNFα as efficiently as conventional drugs used for targeting this pro-inflammatory molecule. Modularity is a key strength of our system. The regulator can be easily swapped, allowing the detection of different biomarkers^27,34^. The cargo (nanobody in our case) can also be replaced in a straightforward manner with other therapeutic proteins, such as small peptides and colonization-increasing factors.

Although mathematical models regarding gut colonization are available^71–75^, they are mostly focused on host-pathogen interactions and not on the colonization-sensing-delivery process from engineered probiotics. Thus, we also developed a simplified yet insightful modelling framework to investigate relevant parameters on probiotic engineering and its subsequent colonization in the gut. Specifically, we investigated the interactions between the probiotic bacteria and inflammation sites, focusing on biomarker (NO) concentration detection thresholds, therapeutic molecule production rates (TNFα), and the bacterial response in terms of therapeutic-target interactions (nanobody-TNFα). We observed that approximately 20 bacteria per mm² are sufficient to manage inflammation. Bacterial density estimates place this requirement at a feasible replacement value of 20 out of 10^4^ gut bacteria per mm² ^69,70^. However, at higher TNFα levels, increased bacterial densities are necessary, suggesting a rough doubling of bacterial density for each magnitude increase in TNFα concentration.

The current understanding of nitric oxide concentrations at inflammation sites within the gut across various patient demographics is limited, with most data focused on serum concentrations^36^. It is estimated that a baseline concentration of around 14 µM NO is typically necessary to detect gut inflammation ^36^. However, we anticipate that the luminal NO concentrations in the gut, particularly at sites of active inflammation, are likely to be considerably higher than this threshold. This expectation is based on the fact that NO, with its notably short half-life and rapid diffusion rates within the body^76,77^, would be more concentrated in regions immediately adjacent to inflammation sites. In light of the scarce available data on serum NO concentrations ^36^, our simulations suggest that an optimal concentration for nanobody production in response to NO is approximately 15 μM. We also observed that enhancing bacterial sensitivity to NO beyond this threshold may lead to diminishing returns. Specifically, this could result in the overproduction of nanobodies in peripheral areas, where they might not contribute effectively to inflammation reduction.

When comparing the impact of the bacterial number on nanobody production per bacterium through simulations, our results indicate that the number of bacteria plays a more critical role than the number of nanobodies produced by each bacterium. This is likely due to the localized distribution of nanobodies, suggesting that a broader gut coverage by bacteria is more effective than increasing the production rate of nanobodies per bacterium. This balance is crucial to avoid the production of excess nanobodies, which could lead to potential side effects and metabolic burden on the bacterial host^78–80^. We highlight that a mathematical model is an oversimplification of reality and does not capture many complexities of the in vivo environment. Future developments in our modelling approach should include important variables such as the consequences of gene expression noise (heterogeneity in gene expression)^81,82^, the reevaluation of the assumptions regarding gut geometry, an enhancement of the diffusion model to encompass three dimensions and the consequences of microenvironmental gut conditions on bacterial growth^83^. Moreover, conducting in vivo studies of the treatment will be instrumental in refining the model as this iterative process of model refinement is essential for advancing our understanding of engineered probiotics^84,85^.

The experimental characterization of our NorR-based circuit revealed that the NO detection threshold in our constructs was higher than the one reported in the original study where this circuit was designed ^38^. This discrepancy could stem from several factors. Firstly, the plasmid used in our experiments differed from that in the referenced study ^38^, and we were unable to access the complete sequences of their constructs, which may have influenced our results. Additionally, our experiments were conducted under aerobic conditions. Previous research has shown that anaerobic environments, akin to the gut’s natural state, can decrease the NO detection threshold of the NorR system by at least five-fold, due to interactions between oxygen and the iron center of NorR ^49^. Consequently, while our sensor system in EcN has been characterized and improved under aerobic conditions, there is substantial potential to enhance its sensitivity to lower, more physiologically relevant NO concentrations. Future studies could achieve this through advanced protein and promoter engineering techniques (e.g., directed evolution and combinatorial designs coupled with fluorescence-based screening methods ^86–89^) and by transitioning to anaerobic assays.

We highlight that although our results support the potential of our system for biotherapeutic applications, the transition from test tubes to translational applications faces many challenges^90–93^, from consistent therapeutic delivery methods to the long-term maintenance of engineered bacteria in the gut. The stable colonization of engineered probiotics in the gut can be negatively impacted by metabolic burden—the allocation of resources towards the engineered system— which can hinder bacterial growth in the complex microbiome environment^79^. Moreover, evolutionary changes might disrupt circuit functionality over short time periods ^94^. The heterogeneity in bacterial expression due to background genetic mutations or expression noise might also lead to variability in treatment efficacy^94^. Additionally, the interactions between the host immune system and engineered probiotics require thorough investigation to ensure long-term efficacy and safety ^95,96^. To address some of these challenges, strategies such as integrating the genetic circuit into the genome can enhance the stability and robustness of the device’s functionality^78,94^. Combining whole-cell and host-microbiome metabolic models with in vivo assays of viability and prevalence of engineered probiotics is also important for predicting the long-term maintenance of such systems ^93,97–101^. Moreover, incorporating antibiotic resistance-free plasmids ^102^ and containment modules ^93,103^ is important to prevent the unintended spread of engineered bacteria and antibiotic resistance genes.

Despite the aforementioned challenges, Synthetic Biology is rapidly transitioning from laboratory experiments to tangible, real-world applications^104–106^. In 2019, ZBiotics Company, USA, pioneered this field by being the first to produce and sell genetically engineered probiotic products, marking the beginning of a burgeoning industry. In a recent notable study, researchers developed a novel system within EcN (PROT3EcT) and validated it in an animal model ^33^. They demonstrated effective colonization of the mouse gut with constitutive production of nanobodies targeting TNFα, resulting in localized inflammation mitigation. Although our system employs different components—specifically, a biomarker-dependent sensing module, distinct nanobodies, and an alternate secretion system—their results are highly encouraging, suggesting the potential functionality of our system in animal models. In this rapidly progressing landscape, our study focused on providing new parts, a new modular system, and a mathematical framework to expand EcN’s Synthetic Biology toolbox and support ongoing efforts in the probiotic engineering community.

## MATERIALS AND METHODS

### Media and buffers

M9 medium is advantageous due to its low cost, low auto-fluorescence (when excited at 488 nm), and low absorbance. We used M9 medium, supplemented with specific amino acids or other metabolites (such as thiamine or casamino acids), for experiments measuring sfGFP fluorescence to ensure minimal auto-fluorescence and absorbance of the samples. To prepare a 50 mL volume of M9 medium, we added the reagents in the following order: 10 mL of M9 salt (5x), 100 µL of MgSO4 (1M), 50 µL of CaCl2 (0.1M), 1.5 mL of Cas Aa (2%), and 1 mL of Glucose (20%), then added water to reach a final volume of 50 mL. If necessary, we supplemented M9 medium with the appropriate antibiotic at a 1:1000 ratio. We conducted all preparation steps under sterile conditions.

### Plate reader fluorescence assay

To measure the activity of all constructs, we transformed plasmids into *E. coli Nissle 1917*. We grew freshly plated single colonies in LB medium supplemented with ampicillin and incubated cultures at 37°C and 220 rpm. On the day of the assay, we spun down the bacteria from the overnight cultures, resuspended them in M9 medium supplemented with ampicillin (M9-Amp) and diluted cultures to OD_600_=0.5. We then assayed the cultures (20 μL) in a 96-well microplate with 170 μL of M9-Amp and 10 μL of the different compounds tested. We used five different concentrations (8 µM, 31 µM, 125 µM, 500 µM and 2000 µM) of the NO donor diethylenetriamine/nitric oxide (DETA/NO) diluted in ddH2O as the inducer. We quantified cell growth (OD_600_) and sfGFP fluorescence using a Tecan Spark 10M plate reader. We calculated the responsiveness of the genetic circuit as arbitrary units using the ratio between fluorescence levels and the optical density at 600 nm (reported as sfGFP/OD_600_) after background correction. As a control for the inducer, we also measured all constructs in the absence of DETA/NO. As a control for cellular autofluorescence background, we also assayed *E. coli Nissle 1917* transformed with the same plasmid but without a promoter to drive sfGFP expression. We measured fluorescence and absorbance at 10-minute intervals for 16 hours at 37 °C and under constant shaking (orbital shaking, 0.1mm orbital averaging). We performed all experiments in technical and biological triplicates. We processed raw data using an ad hoc R script (https://www.r-project.org/).

### Flow Cytometry Analysis

We conducted a high-throughput single-cell analysis of bacteria containing variants of the NO detection module and a negative control plasmid (promoterless *sfGFP*) as follows: first, we selected single colonies of the transformed strain (*EcN*) and cultivated them overnight in LB medium supplemented with ampicillin at 37 °C and 220 rpm. Next, we diluted overnight-grown cells in a ratio of 1:10 in fresh LB and grew them overnight at 37 °C and 220 rpm with different concentrations of the DETA/NO inducer (0 mM,1 mM,1.5 mM, 2 mM). We diluted overnight-grown cells in a ratio of 1:100 in 1mL of filtered cold Dulbecco’s PBS (Sigma-Aldrich #D8537) in 15 mL FACS tubes and immediately stored them on ice to halt metabolic processes.

We set measurements on a BD FACSCantoII machine with the BD FACSDiva 6.1.3 Software after calibration with both CS&T IVD beads and Rainbow Calibration beads (8 peaks, 107/mL, 3.0-3.4 µm, RCP-30-5A) for conversion of arbitrary fluorescence units into MEFL. Excitation and emission filters utilized were 488nm and 530/30 nm, respectively. We adjusted side-scatter (SSC) and forward-scatter (FSC) PMT voltages using bacteria from the negative control, until the distribution of each parameter was centered on the scale. We adjusted FITC/GFP PMT voltage using bacteria from the positive control until the upper edge of the “bell curve” from the fluorescent population was one order of magnitude below the upper end of the scale. We acquired a total of 50,000 events for each biological triplicate and washed cells with PBS before measuring when the bacterial density was too high to avoid the formation of aggregates.

### Nanobodies purification

We transformed *E. coli MC1061* with the pSBinit plasmids containing our 9 different nanobody candidates. The expression vector contains an FX cloning site to insert our different nanobody fragments ordered. The C-terminal myc and 6xHis-tags are included on the plasmid backbone and automatically added in case of a successful FX cloning (Figure 2a). We grew the cells in 600 ml liquid cultures (1:1000 dilution of antibiotic) at 37°C until an OD of 0.4 to 0.7 was reached. We then induced the expression by the addition of 0.02% L-arabinose and allowed bacteria to express the nanobodies for 16 hours at 22°C. We spun down cells at 4’500 rpm for 15 min at 4°C and transferred the resulting supernatant to a bottle with 20 nM imidazole pH 7.5. To extract the nanobodies from the solution, we performed a batch binding using 5 ml Ni-NTA resin for 2 hours while shaking. We poured the resin into gravity flow columns and washed them with TBS pH 7.5, 30 mM imidazole. We eluted nanobodies with 10 ml TBS pH 7.5 and 30 mM imidazole and collected them into fractions, which we measured with the NanoDrop spectrophotometer (Thermo Fisher Scientific). We pooled the fractions with low concentrations and further concentrated them using concentration columns (spun at 2’500 g in 10 kDa concentrators). Lastly, we loaded the purified nanobody candidates on the Sepax in TBS (pH 7.5).

### Cultivation of THP-1 non-adherent human monocytes

We substituted growth medium (RPMI 1640, Gibco) with 10% fetal bovine serum (FBS) and stored at 4°C. We maintained cell densities between 0.1 and 1.0 x 10^6^ cells/ml, splitting them at a ratio of 1:2 or 1:3 (approximately 0.5 x 10^6^ cells/ml) every 3 to 4 days. THP-1 cells display a doubling time of roughly 35-50 hours. During splitting, we transferred cells into 50ml falcon tubes and centrifuged them at 1700 rpm for 5 minutes. We removed the supernatant and resuspended cells in fresh media. After counting the cells, we seeded them at the optimal density.

### Cell Assay

Before the beginning of the actual cell assay, we centrifuged THP-1 cells and resuspended them in starvation media (RPMI 1640 without FBS) and seeded at a density of 1 *x* 10^6^ cells / ml in a 96-well plate (final volume: 200 µl). We then incubated cells for 24 hours at 37°C with 5% CO2, according to the cell specific cultivation protocol.

The next day, we prepared a TNFα dilution series (100 ng/ml, 50 ng/ml, 10 ng/ml, 5 ng/ml, 1 ng/ml, 0.5 ng/ml, 0.1 ng/ml) and kept them on ice. Additionally, we diluted the nanobodies to a final concentration of approximately 100 nM and stored them on ice. After starving the cells for 24 hours, we added the diluted nanobodies to the well plate and gently shook the plate before incubating it for 30 minutes. Afterwards, we stimulated the cells with rTNFα and incubated them for 24 hours at 37°C with 5% CO2. We then harvested the cells, transferred them to Eppendorf tubes, and centrifuged them at 3.5g for 10 minutes at 4°C. After removing the supernatant, we froze the cell pellet with liquid nitrogen and stored it at −80°C for further quantitative RT-qPCR analysis.

### Induction of Nanobody Production and Secretion

We inoculated successfully double-transformed bacteria in 5 ml precultures with a 1:1000 antibiotic dilution and incubated them at 37°C overnight while shaking at 120 r.p.m. The next day, we transferred the cells to 10 ml TB with a 1:1000 antibiotic dilution and grew them at 37°C while shaking until an OD600 of approximately 0.6 was reached. To induce secretion, we added either L-arabinose (final concentration: 0.02%) or diethylenetriamine/nitric oxide (DETA/NO) (testing different concentrations), depending on the transformed cells and their nanobody plasmid. We incubated the cultures at 37°C overnight to allow them to express and secrete nanobodies. We then spun down the cells and collected 2 ml of supernatant for testing via Western Blot or ELISA.

If we needed to test the cell lysate, we first resuspended the cells in TBS and transferred them to a screw-lid microcentrifuge tube. We added one PCR tube of glass beads and lysed the cells using the maxiprep machine at 4 m/s for 20 seconds. We placed the cells on ice for 5 minutes for recovery. We then repeated the shaking process twice, with 5-minute rest intervals in between.

### ELISA

The night before the experiment, we coated a 96-well Nunc Maxicrop immunoplate with 100 µl of protein A solution (1:1000 dilution in PBS) in each well, sealed the plate, and incubated it at 4°C overnight. Before starting the experiment, we freshly prepared the buffers according to the following specifications for ELISA: Tris-buffered saline (TBS) at 1x concentration; TBS-BSA, which is TBS supplemented with 0.5% Bovine Serum Albumin (BSA, weight/volume); TBS-D, consisting of TBS supplemented with a detergent of choice at an amount equivalent to three times the Critical Micelle Concentration (CMC) of the chosen detergent; and TBS-BSA-D, combining TBS with both 0.5% BSA and 0.1% of the chosen detergent (weight/volume). We washed each well with 250 µl TBS and then blocked them with 250 µl TBS-BSA for 30 minutes. We washed the plate three times with 250 µl TBS per well. Then, we added 100 µl of 1:2000 diluted monoclonal anti-c-myc antibody (diluted in TBS-BSA-D) to each well and incubated for 20 minutes. We washed the plate three times with 250 µl TBS-D and added samples diluted in TBS-BSA-D (20 µl in 80 µl solvent for supernatant or periplasmic extraction, or approximately 50 nM for purified nanobodies). We washed the plate three times with 250 µl TBS-D, then added 100 µl of 50 nM biotinylated TNFα in TBS-BSA-D and incubated for 20 minutes. We washed the plate three times with 250 µl TBS-D before adding 100 µl of 1:5000 diluted streptavidin-peroxidase polymer solutions (diluted in TBS-BSA-D) and incubating for 20 minutes. After washing the plate three times with 250 µl TBS-D, we added 100 µl of ELISA developing buffer and incubated until individual wells turned blue, which took between 5 to 15 minutes. We then measured the absorbance at 650 nm using a plate reader. ELISA signals as small as 1.5-fold above the background can indicate a high-affinity binder.

## ASSOCIATED CONTENT

**Supplementary Information** is linked to the online version of the paper

## Acknowledgements

We extend our gratitude to the UZurich 2022 iGEM team for their collective efforts in developing this study. Special thanks to Timothy Kurrer for his support throughout the competition, and to Yen Way Isabell Trinh and Christian Andres Ramos Uria for their insights on our mathematical model. We also thank the UZH flow cytometry facility for their technical support and the gastroenterologists from IBD Net for their feedback. Lastly, we appreciate Professors Markus Seeger from UZH for welcoming us in his group, and Richard Murray from Caltech for sharing his expertise and protocols related to *Escherichia coli Nissle*.

## Author contributions

All authors conceived the study and designed experiments.

**L.B, J.M.**, **M.M.**, **K.J.** and **C.A.W.** carried out NO-induced fluorescent reporter assays, analyzed data, and generated figures.

**G.A.**, **N.W.**, **F.A.** and **J.E.** carried out nanobody production, purification and activity assays, analyzed data, and generated figures.

**G.A.**, **N.W.**, **F.A.** and **J.E.** carried out NO-induced anti-TNF nanobodies secretion assays, analyzed data, and generated figures.

**A.M.** and **A.S** carried out modelling, simulations, and generated figures.

**N. W.**, **M. C.**, **A. S.**, **G. A.** and **M. M.**, contributed with the writing and editing of the paper.

**C.A.W.** wrote the final version of the paper.

## Data availability

The data generated in this study will be deposited in a public repository upon manuscript acceptance.

## Computer code

The computer code will be shared in a public repository upon manuscript acceptance.

## Competing interests

The authors declare no competing interest.

## Supporting Information

The general methods employed in this study include PCR (Section 1.1), real-time quantitative PCR (Section 1.2), preparation of calcium-competent EcN (Section 1.3), heat-shock transformation of calcium-competent EcN (Section 1.4), Gibson assembly (Section 1.5), preparation of electrocompetent EcN (Section 1.6), electroporation (Section 1.7), and Western Blot (Section 1.8). The plasmid design and construction cover secretion plasmid design (Section 2.1), NO-sensing plasmid design (Section 2.2), and plasmid cloning (Section 2.3). Model supplementary methods include assumptions and parameters (Section 3.1), number of inflammatory sites (Section 3.2), number of bacteria (Section 3.3), emission coefficients (Section 3.4), diffusion coefficients (Section 3.5), emission dynamics (Section 3.6), and diffusion dynamics (Section 3.7). Supplementary tables list oligonucleotides used (Table S1). Supplementary figures feature the plasmid map of the negative control (Figure S1), the plasmid map of the engineered nitric oxide sensor construct piGEM2 (ß-1) (Figure S2), the plasmid map of the engineered nitric oxide sensor construct piGEM3 (WT) (Figure S3), the effect of DETA/NO on cellular growth (Figure S4), the impact of removing the plasmid-expressed NorR on NO sensitivity and response strength (Figure S5), the plasmid map of the nanobody purification plasmid (Figure S6), the plasmid map of the arabinose-induced nanobody expression plasmid (Figure S7), the plasmid map of the secretion system plasmid (Figure S8), the plasmid map of the NO-induced nanobody expression plasmid (ß-2) (Figure S9), the purification of monovalent and bivalent anti-TNFα nanobodies from E. coli MC1061 (Figure S10), the comparison of over day to overnight arabinose-induced nanobody secretion in E. coli MC1061 (Figure S11), the arabinose-induced anti-TNFα nanobody production in E. coli MC1061 (Figure S12), the arabinose-induced anti-TNFα nanobody production in E. coli Nissle 1917 (Figure S13), the analysis of ELISA comparing the binding capabilities of purified and secreted monovalent and bivalent anti-TNFα nanobodies in E. coli Nissle 1917 and E. coli MC1061 (Figure S14), the NO-induced monovalent anti-TNFα nanobody expression in E. coli Nissle 1917 (Figure S15), and a visual comparison between diffusion models (Figure S16). (PDF).

## Funding

This study was supported by multiple funding sources. Academic funding was provided by the University of Zurich through the UZH Alumni Science, the Faculty of Science (MNF), the Faculty of Medicine, the Vice President Research, and the President’s Services. Private sponsorship was provided by Pierre Fabre, Microsynth, the Swiss Academy of Sciences (SCNAT), and Promega. Additionally, this publication was funded by the University of Zurich and the Consortium of Swiss Academic Libraries, making it available as open access.

## Notes

### Competing Interest Statement

The authors have declared no competing interest.

