## Supplementary Material for "Engineering a novel probiotic toolkit in *Escherichia coli Nissle1917* for sensing and mitigating gut inflammatory diseases"

**INDEX**


1. **General Methods**
   - **1.1.** PCR
   - **1.2.** Real time quantitative PCR
   - **1.3.** Preparation of Calcium competent EcN
   - **1.4.** Heat-shock transformation of calcium competent EcN
   - **1.5.** Gibson assembly
   - **1.6.** Preparation of electrocompetent EcN
   - **1.7.** Electroporation
   - **1.8.** Western Blot

2. **Plasmid design and construction**
   - **2.1.** Secretion plasmid design
   - **2.2.** NO-sensing plasmids design
   - **2.3.** Plasmid cloning

3. **Model supplementary methods**
   - **3.1.** Assumptions and parameters
   - **3.2.** Number of inflammatory sites
   - **3.3.** Number of bacteria
   - **3.4.** Emission coefficients
   - **3.5.** Diffusion Coefficients
   - **3.6.** Emission Dynamics
   - **3.7.** Diffusion Dynamics

4. **Supplementary Tables**
   - **4.1.** Table S1. List of oligos used in this study

5. **Supplementary Figures**
   - **5.1. Supplementary figure S1.** Plasmid map of the negative control
   - **5.2. Supplementary figure S2.** Plasmid map of the engineered nitric oxide sensor piGEM2 (ß-1)
   - **5.3. Supplementary figure S3.** Plasmid map of the engineered nitric oxide sensor construct piGEM3 (WT)
   - **5.4. Supplementary figure S4.** DETA/NO has little effect on cellular growth
   - **5.5. Supplementary figure S5.** The removal of the plasmid-expressed NorR reduces sensitivity and response strength to NO
   - **5.6. Supplementary figure S6.** Plasmid map of the nanobody purification plasmid
   - **5.7. Supplementary figure S7.** Plasmid map of the arabinose-induced nanobody expression plasmid
   - **5.8.Supplementary figure S8.** Purification of monovalent and bivalent anti-TNFα nanobodies from *E. coli MC1061*
   - **5.9. Supplementary figure S9.** Comparison of over day to overnight arabinose-induced nanobody secretion in *E. coli MC1061*
   - **5.10. Supplementary figure S10.** Comparison of TNFα binding capacity between monovalent and bivalent nanobodies
   - **5.11. Supplementary figure S11.** Plasmid map of the secretion system plasmid
   - **5.12. Supplementary figure S12.** Arabinose-induced anti-TNFα nanobody secretion in *E. coli MC1061*

5.13. **Supplementary figure S13.** Arabinose-induced anti-TNFα nanobody secretion in *E. coli Nissle 1917*
5.14. **Supplementary figure S14.** Analysis of ELISA comparing the binding capabilities of purified and secreted monovalent and bivalent anti-TNFα nanobodies in *E. coli Nissle 1917* and *E. coli MC1061*
5.15. **Supplementary figure S15.** Plasmid map of the NO-induced nanobody expression system (ß-2)
5.16. **Supplementary figure S16.** NO-induced monovalent anti-TNFα nanobody secretion and function in *E. coli Nissle 1917*
5.17. **Supplementary figure S17.** Visual comparison between diffusion models

**6. References**

### 1. General Methods

#### 1.1. PCR

To amplify inserts, PCR was performed with Phusion Hotstart polymerase. Amounts of reactants for one aliquot: 31 µL ddH2O, 10 µL Hi-Fi Phusion buffer (5x), 2.5 µL of each, forward and reverse primer (10 µM), 1 µL DMSO, 1 µL MgCl2 (50 mM), 1 µL dNTP (10 mM), 0.5 µL of template (1 µL for low-concentrated templates), 0.5 µL Phusion polymerase. We performed PCR with the following program: 98 °C/3 min; 25 cycles of 98 °C/30 s, 50 °C/30 s, 72 °C/0.5 min/kb; 72 °C/10 min. Following PCR, the fragments were separated by electrophoresis on a 1% agarose-gel and subsequent gel extraction.

#### 1.2. Real time quantitative PCR

The frozen cell pellets were thawed and lysed in order to isolate the RNA using Maxwell RCS (promega) according to the company's protocol. Then reverse transcription was performed using the TaqMan™ Reverse Transcription Reagents kit (Thermo Fisher N8080234) to obtain the cDNA of the transcriptome of the cells. Reverse transcription was performed in a thermocycler and kept at 4°C until further processed. The obtained cDNA was diluted to 25ng/µl and prepared for the qPCR. ABI Fast Polymerase mix (Applied Biosystems) was used and primers for IL-1ß (gene of interest) were added together with primers for GAP-DH serving as the house-keeping gene. Samples were pipetted as triplicates in a 384-well plate and qPCR analysis was performed with the QuantStudio 6 Real-Time PCR system (Thermo Fisher Scientific).

#### 1.3. Preparation of calcium competent EcN

*E.Coli Nissle 1917* bacteria were obtained from Mutaflor® (Herdecke, Germany) and cultured overnight in LB medium at 37°C, 220 rpm. This original culture was diluted 1:10 and 1:100 on the following day and 100 µL of each dilution were plated on agar-plates (overnight, 37°C, 220 rpm). Competent bacteria were made from a single colony from one of the plates (where easier to pick one) following an in-house established protocol.

#### 1.4. Heat-shock transformation of calcium competent EcN

10-100 ng of the vector were added to ice-cold 50 µL aliquots of chemically competent cells and incubated on ice for 30 min/5 min for miniprepped plasmids. Following this, cells were heat-shocked for 45-50 s at 42°C and placed back on ice for 5 min. 350 µL LB medium was added and tubes were incubated at 37°C, 200-300 rpm for 1h/15 min for miniprepped plasmids. Tubes were then spun down for 5 min at 3000g, the supernatant poured away and the pellet resuspended in the remaining liquid. (This step can be skipped for miniprepped plasmids.) From the transformed bacteria, up to 50 µL were plated on small ampicillin-supplemented agar plates (1:1000) and cultivated at 37°C, 220 rpm.

#### 1.5. Gibson assembly

First, an in-house Gibson Master Mix was prepared and stored in 15 µL aliquots at -20°C. Formula for a 1.2 mL Master Mix: 320 µL ISO buffer (5x), 699 µL ddH2O, 160 µL Taq ligase (40U/µL), 20 µL Phusion polymerase (2U/µL), 0.64 µL T5 exonuclease (10U/µL). The linearized backbone and fragments were mixed in a 1:2 ratio into a final volume of 5 µL. This was then added to one aliquot of Gibson Master Mix and incubated for 1h at 50°C. For transforming bacteria with a Gibson assembled product, aliquoted chemically competent cells were thawed on ice for 10 min, then mixed with 5 µL of the freshly made Gibson mix and incubated on ice for 1h. Following this, cells were heat-shocked for 45 sec at 42°C, then placed back on ice for 3 min. 0.5 mL LB medium was added and tubes were incubated for 1h at 37°C, 600 rpm. From the transformed bacteria, 100 µL were plated on small agar plates, supplemented with ampicillin (1:1000), and cultivated at 37°C, 220 rpm.

**1.6. Preparation of electrocompetent EcN**

All tubes and pipettes were prechilled at 4°C or -80°C as appropriate. (Additionally, all flasks were rinsed with $H_2O$ prior to autoclaving in order to remove residual detergents that may remain on glassware from dishwashing. This step may increase competency. Autoclaving with water, which is then discarded, is even better.) EcN was inoculated in 5 ml LB medium and grown overnight at 37°C with rotation. On the next day, 5 ml of overnight cultures were added to 450 ml LB medium and incubated at 37°C with vigorous shaking until the OD 600 nm was between 0.5 and 1.0. This step usually takes about 3 hours. The centrifuge was fast-cooled with the correct rotor at 4°C and cultures were poured into two 225 ml centrifuge tubes. The tubes were placed on ice for 15 minutes. Longer incubation up to 1 hour is possible and may lead to higher competency.

*For the following steps it is important to keep cells cold and remove all the supernatant in each step to remove residual ions.*

The cells were centrifuged for 10 minutes at 2’000g at 4°C. Afterwards, the supernatant was removed and the cell pellets were gently resuspended with 200 ml cold sterile water. Initially, 10 - 20 ml of cold water was used to resuspend the pellet by pipetting and then the rest of the water was added. The cells were centrifuged again for 15 minutes at 2’000g at 4°C. The supernatant was removed and the pellets were resuspended with 200 ml cold sterile water. The cell suspensions were held on ice for 30 minutes before they were centrifuged for the third time for 15 minutes at 2'000g at 4°C. The supernatant was removed and the cell pellets were resuspended with 25 ml cold 10% glycerol. The mixture can be optionally transferred to a 50 ml conical tube. The cells were placed on ice for 30 minutes. Afterwards, a next centrifugation step for 15 minutes at 1'500g and 4°C was performed and the supernatant was removed. 500 µl of 10% glycerol was added to the pellets and the cells were resuspended in a final volume of approximately 1 ml. 50 µl aliquots were prepared (tubes on ice) and the cell suspension was shock frozen in a dry ice and ethanol bath. The aliquots were then stored at -80°C.

**1.7. Electroporation**

1.5 ml reaction tubes were prepared containing 100 ng of each plasmid DNA (correct Nb & SS plasmid names according to list). The electroporation cuvettes (electroporation cuvettes plus, model no. 610, 1 mm) and reaction tubes containing the DNA were placed on ice. Electrocompetent *E. coli Nissle 1917* cells were thawed on ice for about 10 minutes and 40 µl of EcN was added to the reaction tubes and mixed well by flicking the tubes gently. The mixture was then transferred to a chilled microcentrifuge tube. The cell / DNA suspension was carefully transferred into a chilled cuvette without introducing bubbles. It is important that the cells deposit across the bottom of the cuvette. The electroporation (Gene Pulser Xcell electroporation system) was then performed using the following conditions: 1800 V, 600 Ω, and 10 µF. The typical time constant is approximately 4 milliseconds. After the electroporation, 1 ml of LB medium was immediately added to the cuvette and gently mixed up and down twice before the cells were transferred to a new 1.5 ml reaction tube. The cells were incubated for 30 minutes while shaking at 37 °C and 160 r.p.m. for recovery. Afterwards, 100 µl of cells were spread onto selective plates, supplemented with ampicillin and chloramphenicol. For liquid cultures, 100 µl cells were added into 5ml selective media, once with normal antibiotics concentrations (5ul Amp, 2.5ul Chlor) and once with half the concentrations (2.5ul Amp, 1.25ul Chlor). The plates and liquid precultures were incubated at 37°C overnight.

**1.8. Western Blot**

To quantify the presence of nanobodies in the supernatant of double-transformed and induced EcN, a western blot was performed. 50 µl of supernatant (or lysate in the case of testing for intracellular nanobodies) were added to 12.5 µl 5x Protein loading dye. 20 µl of the samples were run on a 4 - 20% gradient gel in MOPS buffer for 50 minutes at 140 V. The gel and blotting paper were soaked in transfer buffer (20 mL 100% methanol, 20 mL 10x transfer buffer, 0.2g SDS, 160 mL water). The membrane was first soaked in 100% methanol before placed into the transfer buffer. The assembly of the blot was then performed as following (from top to bottom): Blotting paper - gel - membrane - blotting paper.

The transfer was conducted at 12 V in Trans-blot SD semi dry transfer cell for 0.5 - 1 hour. In the meantime, 800 ml of PBS-T (0.05% Tween 20 added to PBS) and 250 ml of blocking buffer (250 ml PBS-T and 7.5 g BSA) were prepared. After the transfer, the membrane was blocked for 0.5 - 1 hour in blocking buffer while shaking at room temperature. The membrane was then incubated with the primary antibody (5 ml blocking buffer and 1 µl anti-myc antibody). The membrane was placed into a 50 ml falcon tube containing the primary antibody solution and incubated for 0.5 to 1 hour while rotating. The membrane was washed three times with PBS-T for 5 minutes while shaking. The secondary antibody solution was prepared using 25 ml blocking buffer and 1 µl anti-mouse antibody. The membrane was incubated with the secondary antibody for 0.5 to 1 hour while shaking. Afterwards, it was washed three times with PBS-T for 5 minutes while shaking. Imaging was performed with an Image Quant 800. A 1:1 ratio of immobilon western blot HRP substrate peroxidase solution and immobilon western blot HRP substrate luminol reagent were mixed (1 ml per membrane required) in an Eppendorf tube. The developing solution was slowly added to the membrane and bands were imaged (chemiluminescence setting with colorimetric marker for ladder).

To analyze the relative intensities of the bands we used the protocol by Hossein Davarinejad[1] and visualized the data with R.

### 2. Plasmid design and construction

#### 2.1. Secretion plasmid design

All plasmids were designed in Benchling and the sequences for the HlyB and HlyD of the secretion system plasmid as well as the HlyA-tag integrated in the nanobody plasmid were obtained from [2]. TolC is endogenously expressed in *E. coli* strains and is therefore not necessary to be integrated in a plasmid. Generally, all promoter, RBS, and double terminator sequences were obtained from the corresponding iGEM parts registry. The arabinose-inducible system consisting of the pBad promoter and araC, as well as the myc-tag were adapted from the pSBinit [3] plasmid (addgene #110100).

#### 2.2. NO-sensing plasmids design

All plasmids were designed in Benchling and the sequences for pNorVβ, sfGFP and NorR were obtained from Chen XJ et al. [4]. NorR was further optimized to avoid repetitive sequences. Generally, RBS and double terminator sequences were obtained from the corresponding iGEM parts registry. The sequence for the wild-type pNorV was obtained from previous iGEM work ( http://parts.igem.org/Part:BBa_K2116002 ).

The different nanobody candidates were ordered as fragments from IDT. The amino acid sequences of the nanobodies used in this study were taken from the patent of Karen Silence et al[5] (Int. Publication Number: WO 2004/041862 A2) and converted to their corresponding DNA sequences using the Expasy software.

Codon optimization for *E. coli* was performed on all plasmids and DNA fragments using the integrated codon optimization tool offered by Twist Bioscience.

#### 2.3 Plasmid cloning

Adjusting the number of RBS upstream of GFP as well as combining the NO-sensor with the nanobody (Nb1) were performed by Gibson assembly. For this purpose, fragments were ordered from IDT (for the RBS) or linearized from a miniprepped plasmid vector (for the nanobody) and mixed with the miniprepped linearized backbone following the protocol described earlier. Pure linearized fragments and backbones were obtained by gel extraction.

### 3. Model supplementary methods

The gut surface section is constructed as a square matrix with N rows and N columns with each entry representing a 1µm$^3$ volume and the entire grid representing an area of 1mm$^2$. Some grid areas are randomly assigned the status "inflamed," and start producing NO and TNFα. After the initial setups, TNFα levels decide whether the status “inflamed” is maintained. If TNFα levels drop below a certain threshold, the status switches to “uninflamed”. *E. coli* bacteria are randomly distributed across the grid and occupy a single instance of our grid as they have a rough volume of around 1µm$^3$. If the grid cell of a bacteria reaches an NO concentration above their sensing threshold, they produce nanobodies in their grid element. All particles are measured in mol/µm$^3$.

The particles (NO, TNFα, and nanobodies) are subject to diffusion and decay over time. If concentrations of nanobodies and TNFα overlap in the same 1µm$^3$, we assume that they will bind and cancel each other out in a 3:1 nanobodies:TNFα ratio, as we target three possible binding sites.

Our model follows a cycle of operations comprising four steps in the following order: 1) particle production, 2) particle diffusion and decay, 3) nanobody and TNFα binding and canceling, and 4) data collection or plotting.

#### 3.1. Assumptions and parameters

We made the following simplifications and assumptions in our model: Bacteria attach to the gut surface and remain static without dying/turnover, NO sensing and nanobody production is immediate, without any time lag, inflammation sites can only shrink and not expand, and the compounds only interact with themselves during diffusion.

**Number of inflammatory sites**

This parameter corresponds to the count of inflammatory sites generated. Given the broad variability among human patients with IBD, the parameter was arbitrarily set to a default of 50 inflammation sites with variable sizes, to represent a broad range of conditions.

**Number of bacteria**

The amount of bacteria that are able to remain in the gut and produce nanobodies is crucial for the efficacy of the treatment, but hard to assess without further studies into the fitness of our engineered bacteria. Studies have estimated the bacterial density in the colon as $10^{11}$ per milliliter of gut content[6]. In our model this equates to a probability of around 0.1 that a grid entry is filled with bacteria. For our simulation we chose a default value such that our treatment will replace around 20 out of an estimated $10^5$ gut bacteria per $mm^2$ [7,8]. This gives a sufficient coverage of the gut based on our simulations.

#### 3.4. Emission coefficients

Each grid element that is part of an inflammation site produces a fixed amount of NO and TNFα. Since there is no data on NO and TNFα concentrations around inflammation sites in the gut, we used the values from the medium concentrations in blood serum samples of IBD patients, with NO concentrations between 14.54 µmol/L and 15.25 µmol/L [9]. We used a default value of 15 µmol/L and changed it into our standard unit to get 1.5 $x$ $10^{-20}$ mol/µ$m^3$. TNFα concentrations in the blood serum of UC patients lie around 8.3 ± 2.5 pg/ml and in CD around 5.4 ± 1.7 pg/ml [10]. We chose a default value of 5.4 pg/ml which results in 3.12 $x$ $10^{-28}$ mol/µ$m^3$ when considering a weight of 17.4kD. As these are rough estimates, a lot of different concentrations have been tested, and do not seem to greatly influence the efficacy.

If a grid element contains bacteria and the concentration of NO is above the sensing threshold of the bacteria, nanobodies are produced. The grid element's nanobody concentration increases by a default concentration of 1.66 $x$ $10^{-21}$ mol/µ$m^3$. The value is extrapolated from the lower bound concentrations produced by an *E.coli* population[11]. However, this concentration is only reached if the simulation were to assume complete colonization of the gut. The actual values could greatly differ and as such have been explored in our model.

We used 2.6 $x$ $10^{-20}$ mol NO/µ$m^3$ as our default sensing threshold, but a recent paper has shown a ten times more sensitive threshold [4], which might be necessary for the treatment. Experiments were made with the assumptions that we could replicate the results and work with a higher sensitivity.

#### 3.5. Diffusion Coefficients

The diffusion coefficients used are 3300 µm² per second for NO [12] and 7.28 µm² per second for TNFα based on proteins of similar size [13,14] for the nanobodies we chose 40 µm² based on the same calculations [15], which is similar to the upper bound for antibodies [16]. The actual diffusion speed, however, is likely to be higher and could further improve the efficacy.

Every particle that leaves its generative environment through diffusion will eventually decay. To simulate this, we enforce half-lives of each particle. We used a 2 seconds half-life from studies in extravascular tissue [17]. For TNFα, we used parameters from a study about the half-life of TNFα from intravenous injections in rats. The researchers found near dose-independent decay of around 30 minutes half-life in the high-dosage conditions [18]. As all nanobodies are structurally similar, we used a half-life estimate of 12 minutes which is the average half-life described in a paper about nanobodies as imaging agents [16]. Some studies have shown that the half-life can be extended up to multiple days [19], which would trade ease of production for a longer lifespan.

#### 3.6. Emission Dynamics

The emission rates of NO and TNFα particles were designed to maintain a constant particle density by compensating for the losses. When increasing the time-scale model, we need to ensure that sufficient particles are introduced to bridge the period where no additional particles are added. To illustrate a transition from a timestep of n seconds to $10 * n$ seconds, consider an experiment involving two buckets of water. In the first trial, the initial bucket contains $e$ liters of water. We transfer a proportion $k$ of the water to a second bucket, and subsequently refill the original bucket to maintain the initial $e$ liters. This process repeats n times, resulting in a final quantity of water $V$ in the second bucket given by equation [1].

$$V = n * k * e$$

[1]

In the second trial, no refilling takes place and instead we increase the initial volume to a value of $e_n$ liters, so that the same amount of water is transferred to the second bucket over n timesteps, even without refilling the bucket. This corresponds to the need to have an equal amount of particles spread out through the diffusion and decay-steps over the same number of time. This achieves the same final volume in the second bucket, even when we transfer proportion $k * n$ times without refilling. Given the decreased water volume on each transfer, the first transfer yields $k * e_n$ liters, followed by $k * (1 - k) * e_n$ liters for the second transfer. The sum of these transfers over n iterations should equal the volume $V$ from the first trial and results in equation [2].

$$V = \sum_{i=0}^{n-1} k * (1-k)^i * e_n$$

[2]

With [1] and [2] we solve for $e_n$, we derive [3]:

$$\frac{n * e}{\sum_{i=0}^{n-1} (1-k)^i} = e_n$$

[3]

The emission values e from [1] are therefore replaced by $e_n$ when scaling the model to higher time-scales.

#### 3.7 Diffusion Dynamics

After the production of particles, they diffuse from their origin. This process can be modeled using the Heat Equation. By discretizing this partial differential equation (where the Mesh Fourier number $F$ corresponds to the product of the diffusion coefficient and the difference in time over the difference in distance), the propagation of particles in one dimension can be simulated utilizing the backward Euler scheme.

For a one-dimensional parameter vector, a two dimensional diffusion matrix $D$ is needed. The principal diagonal of the matrix contains the value $1+2F$, while the two adjacent diagonals contain the value $-F$. As every particle has a unique diffusion coefficient, a dedicated diffusion matrix is required for each particle. Given the concentration within a particular grid element, the new concentrations after one time-step can be calculated by multiplying the parameter vector $v$ with the inverse of the diffusion matrix.

The diffusion process can be extrapolated from one dimension to a two-dimensional parameter concentration space $M$, by multiplying the parameter space as $D^{-1}MD^{-1}$. For a more accurate 2D diffusion simulation, a Crank-Nicolson scheme in combination with the Runge-Kutta scheme could be used. However, this method is significantly more computationally demanding as it requires a diffusion matrix of size $N^2*N^2$ of the initial matrix size. To compare the performance of the two methods, we simulated 30 timesteps using a parameter space of size 100µm² with an arbitrary diffusion coefficient of 20 µm²/s, and a starting concentration of 1 mol/µm³ at index $x$=50 and $y$=50 and evaluated the resulting diffusion patterns. (**see Supplementary figure S17**. The approximation results in practically indistinguishable diffusion patterns for high diffusion coefficients and confirm our choice of diffusion modeling.

To compensate for the discrete emission and diffusion, scaling to larger time-steps needs to be compensated. The parametric representation of our particle matrix is given by the current concentration space $M$. We replaced the emission e with $e_n$ and represent it as the emission matrix $E_n$, where $D$ is our diffusion matrix and $p$ represents our decay parameter. The number of repeats is denoted by $n$. When increasing from time-step $n$ to 10*$n$, a continuous induction of particles can be simulated by diffusing 1/10th of the particles ten times, 1/10th diffuse nine times, and so forth. The product of these diffusion matrices can be calculated at the beginning of the simulation and used as the new diffusion matrix to calculate the diffusion of 10*$n$ timesteps with the same number of matrix multiplications per timesteps. The same applies for the decay of the particles. We can calculate this diffusion matrix, including the decay parameters at the beginning of the simulation in [4]:

$$D_{pn} = \sum_{i=1}^{n} \frac{D^{-i} * \sqrt{p^i}}{n}$$

[4]

The square root of $p$ is taken, as we multiply the diffusion matrix twice in the step update. Before we updated $M$ in every timestep. With the new diffusion matrix $D_{pn}$, $n$ steps of simulation can be calculated in a single step as in [5]:

$$(D^{-1}(M + E) * p * D^{-1})^n = D_{pn}(M + E_n)D_{pn}$$

[5]

To discrete the diffusion accurately, an initial diffusion matrix for 1 millisecond is used to calculate the final diffusion matrices.

### 4. Supplementary Tables

#### 4.1. Table S1. List of oligos used in this study:

| Name | Sequence |
|---|---|
| oiGEM15 (fwd) - +Nb | TAAGCTCTTCGTGGAAAGAGGAGAAAATGAGTTTTAGCGTTGAC |
| oiGEM16 (rev) - +Nb | CTCTTTCCACGAAGAGCTTATTATGCTGATGCTGTCAAAGTTATTG |
| oiGEM17 (fwd) -+Nb | ATGAGTCAAGTCCAATTACAGGAGAGCGGTGGCGGGC |
| oiGEM18 (rev) - 1RBS +Nb | CCTGTAATTGGACTTGACTCATTTTCTCCTCTTTCTAATGAAGAGCC |
| oiGEM20 (rev) - 2RBS+Nb | CCTGTAATTGGACTTGACTCATCATCTAGTATTTCTCCTCTTTGGTTTC |
| oiGEMnoNOR1 (fwd) - remove NorR | CTCTTCGTGGCCAGGCATCAAATAAAACGAAAGGCTCAGTCGAAAG |
| oiGEMnoNOR2 (rev) - remove NorR | GATGCCTGGCCACGAAGAGCTTATTTGTAGAGCTCATCCATGCC |
| oiGEMrbs1 (fwd) - for 3RBS | AGGAGGTTTGGATTCACACAGGAAACCAAAGAGGAGAAATACTAGATGATGAGCAAAGGAGAAGAACTTTTCAC |
| oiGEMrbs2 (rev) - for 3 RBS | CATCTAGTATTTCTCCTCTTTGGTTTCCTGTGTGAATCCAAACCTCCTCTAATGAAGAGCCTAAAAAGATGTCTTGC |
| oiGEMrbs3 (fwd) - for 2 RBS | TTCACACAGGAAACCAAAGAGGAGAAATACTAGATGATGAGCAAAGGAGAAGAACTTTTCAC |
| oiGEMrbs4 (rev) - for 2 RBS | CATCTAGTATTTCTCCTCTTTGGTTTCCTGTGTGAACTAATGAAGAGCCTAAAAAGATGTCTTGC |
| M13 fwd - sequencing | GTAAAACGACGGCCAGT |
| M13 rev - sequencing | GTCATAGCTGTTTCCTG |

### 5. Supplementary Figures

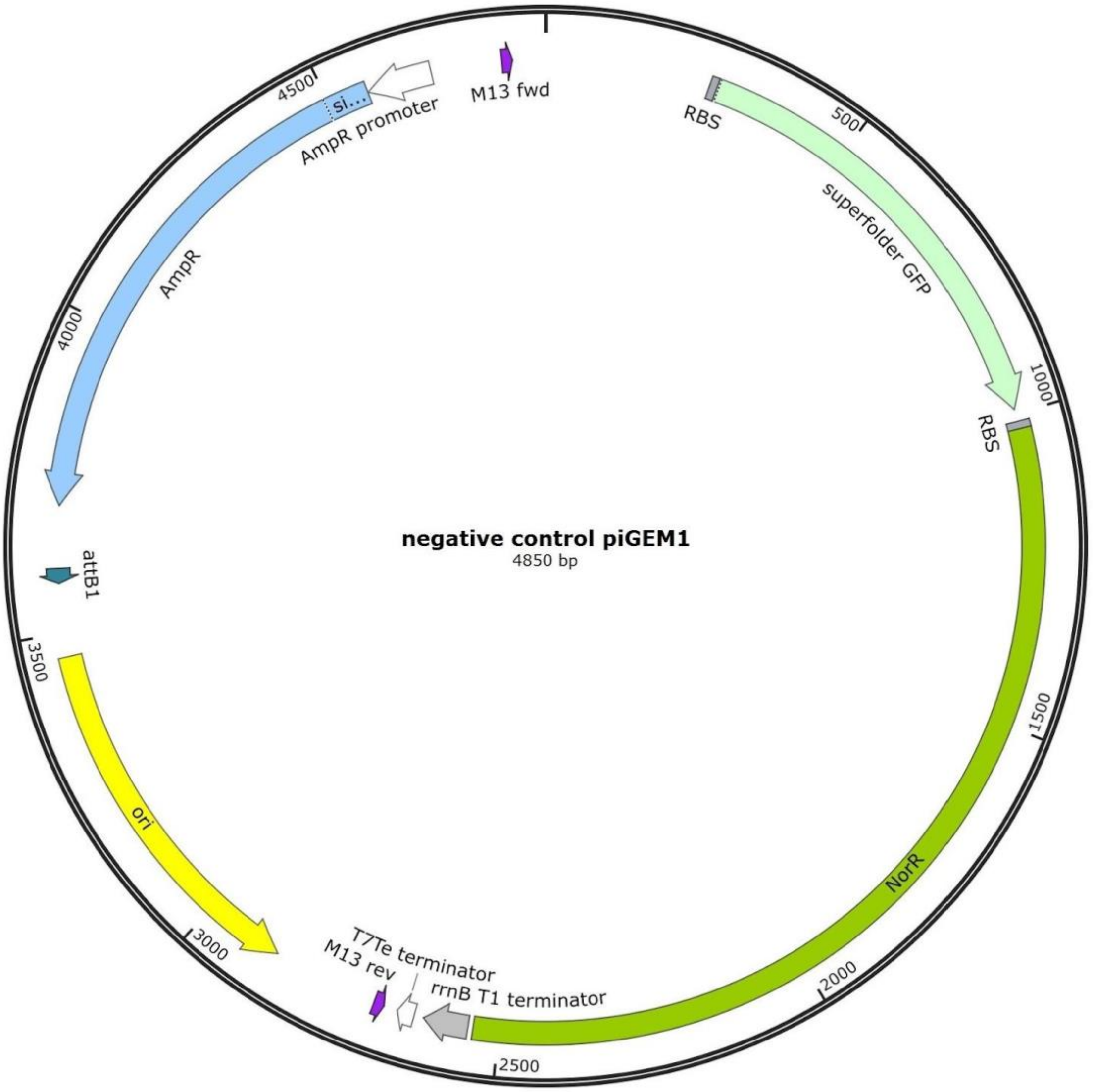


**5.1. Supplementary figure S1. Plasmid map of the negative control.** The negative control plasmid encodes a high copy number origin (colE1), a superfolder GFP, the NorR gene for the positive feedback loop, but no pNorVß promoter. This plasmid was used for the normalization of the plate reader fluorescence assay data to characterize the NO sensor.

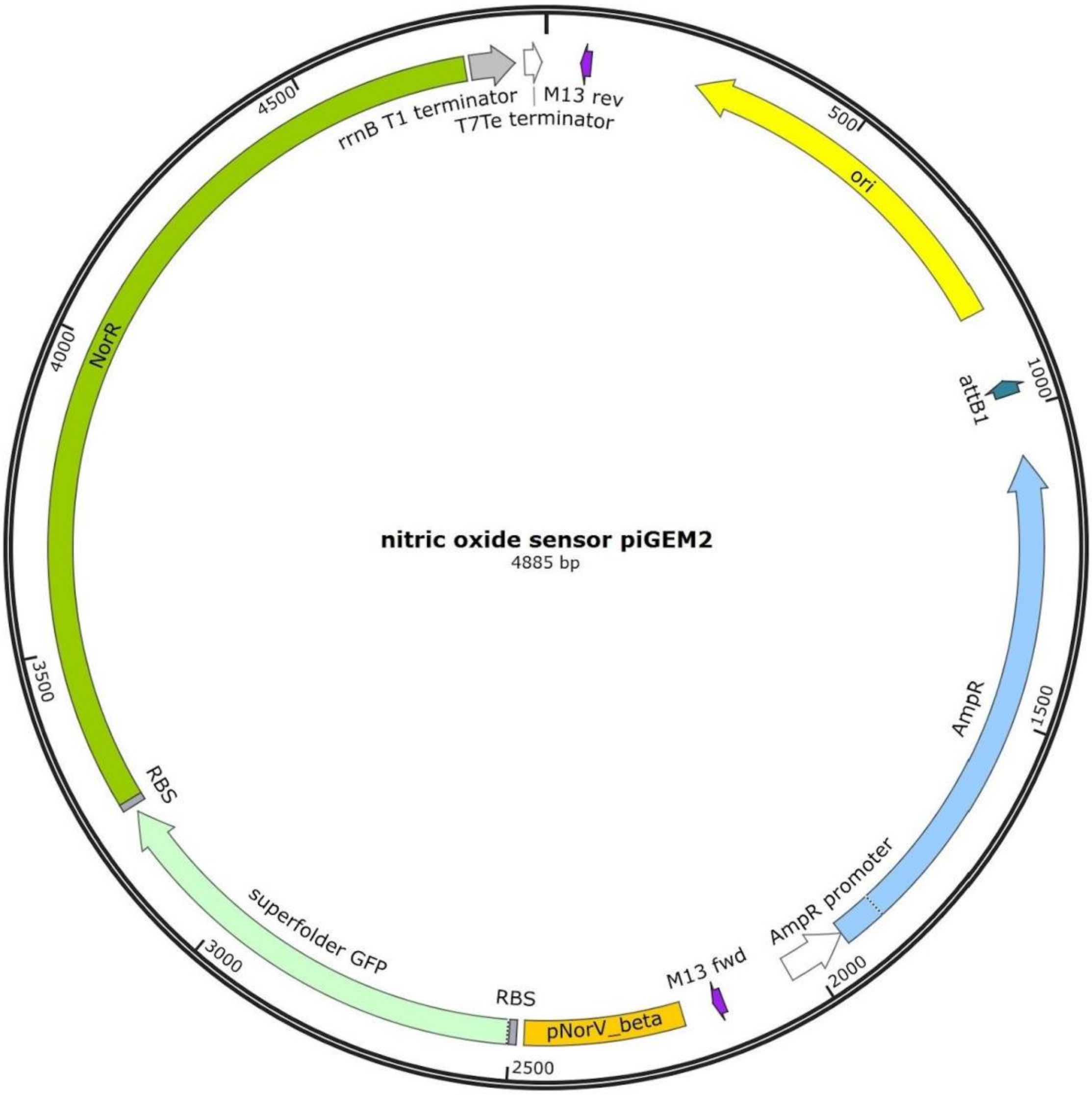


**5.2. Supplementary figure S2. Plasmid map of the engineered nitric oxide sensor piGEM2 (ß-1).** The piGEM2 plasmid encodes a high copy number origin (colE1), a superfolder GFP, the NorR gene for the positive feedback loop, and the pNorVß promoter preceded by 1 RBS. Via Gibson Assembly, this plasmid was further modified to obtain the ß-2 and ß-3 plasmids containing two or 3 RBS. This plasmid was used for the plate reader fluorescence assays, to characterize the NO sensor.

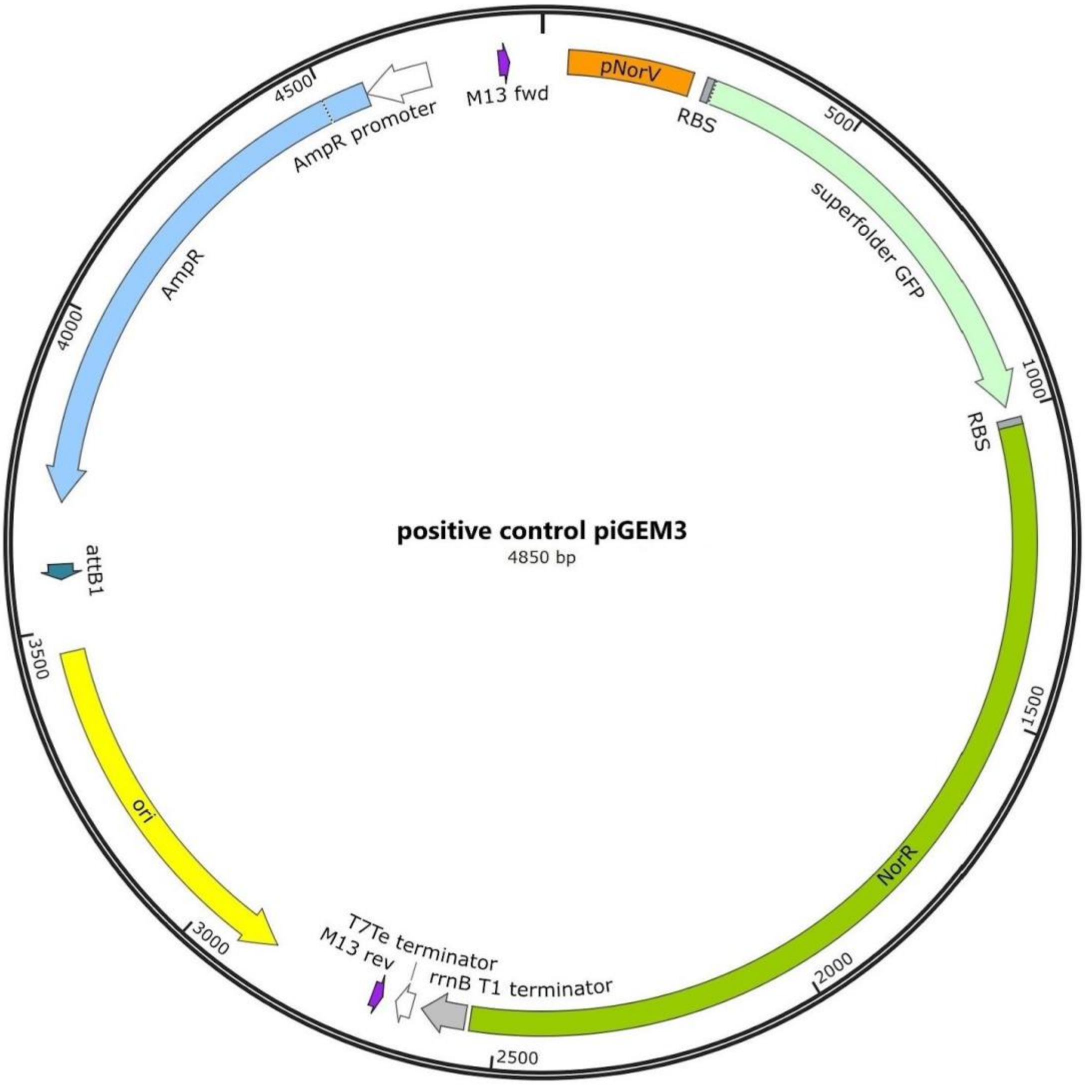


**5.3. Supplementary figure S3. Plasmid map of the engineered nitric oxide sensor piGEM3 (WT).** The piGEM3 plasmid encodes a high copy number origin (colE1), a superfolder GFP, the NorR gene for the positive feedback loop, and the wild-type pNorVß promoter preceded by 1 RBS. This plasmid was used as a positive control for the plate reader fluorescence assays, to characterize the NO sensor.

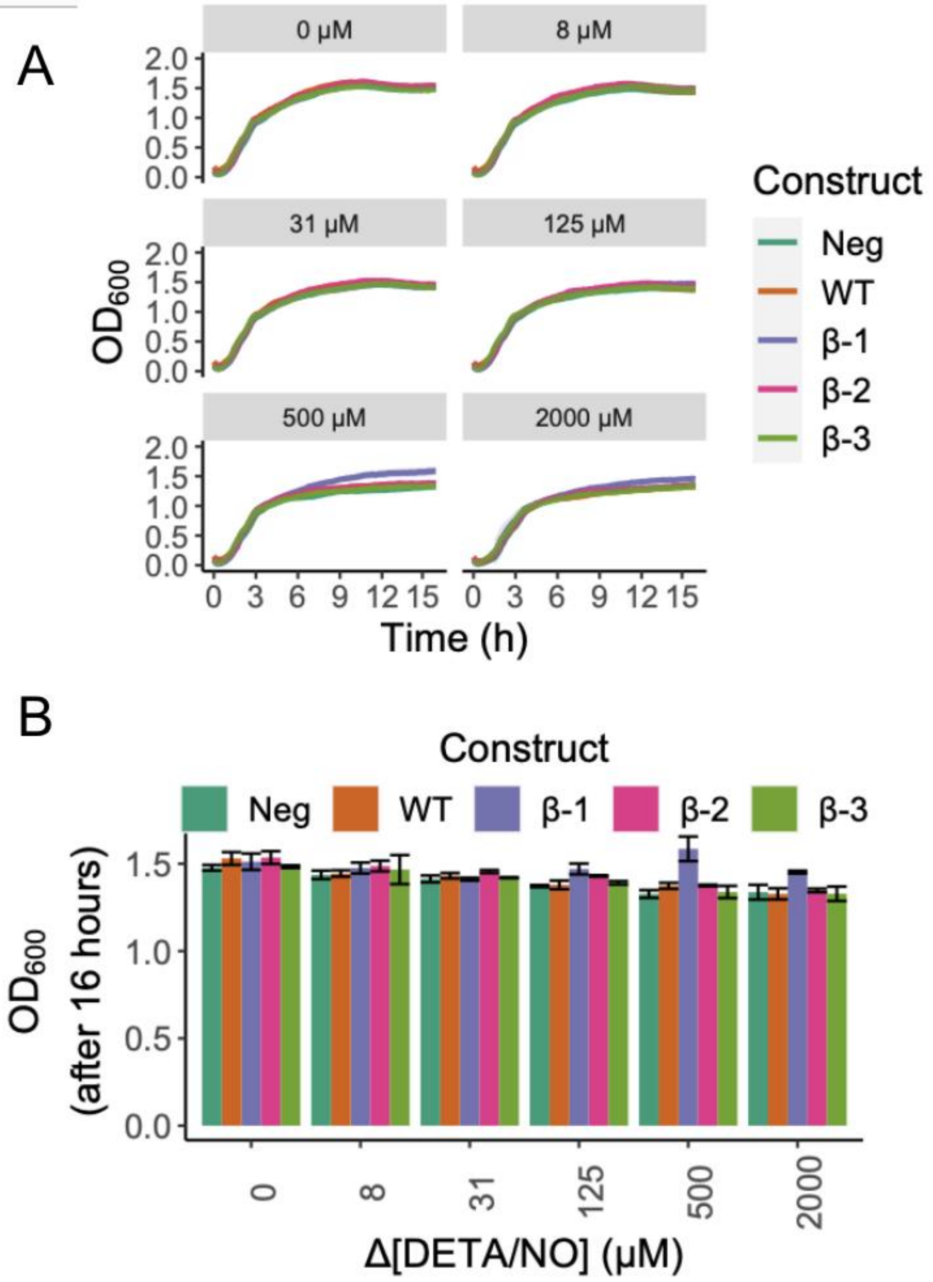


**5.4. Supplementary figure S4. DETA/NO has little effect on cellular growth.** a. Time-lapse growth assay for constructs on different NO concentrations. We have grown each construct for 16 hours (x-axis) on a microplate reader where OD600 was measured every 15 minutes. The y-axis represents OD600 values. Each grid represents a different concentration of DETA/NO in which cells harboring each construct were grown. DETA/NO gradients used were 0, 8, 31,125,500, and 2000μM. Each line color represents a construct. Line shadings represent the standard deviation of our biological replicates (*n=3*). We performed all measurements with both biological and technical triplicates. **b. Endpoint growth measurement for constructs on different NO concentrations.** The bar plots represent the OD600 for each construct for each DETA/NO change of concentration at T = 8h. Error bars represent the standard deviation of our biological replicates (*n=3*). A slight decrease in OD600 values can be observed with incremental NO concentrations due to its cellular toxicity.

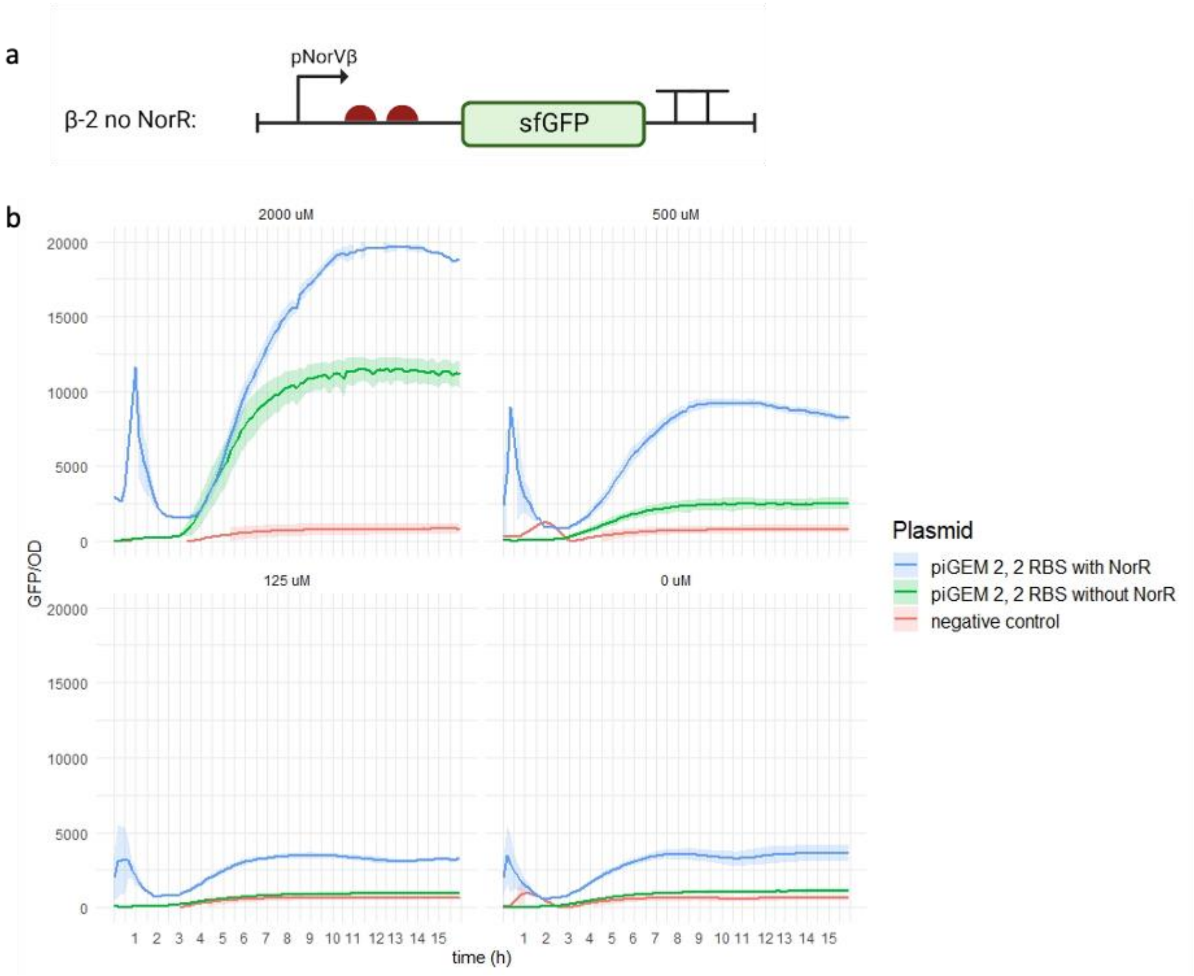


**5.5. Supplementary figure S5. The removal of the plasmid-expressed NorR reduces sensitivity and response strength to NO. a. Schematic representation of β-2 without NorR.** To underline the importance of a positive feedback loop in the sensing module, we also tested the circuit with 2 RBSs after removal of the transcription factor NorR and its corresponding RBS. b. R**esponse of our construct with 2 RBSs +/- NorR to induction with DETA/NO.** The removal of NorR disabled the positive feedback mechanism and did not improve the sensitivity of our construct to nitric oxide (NO).

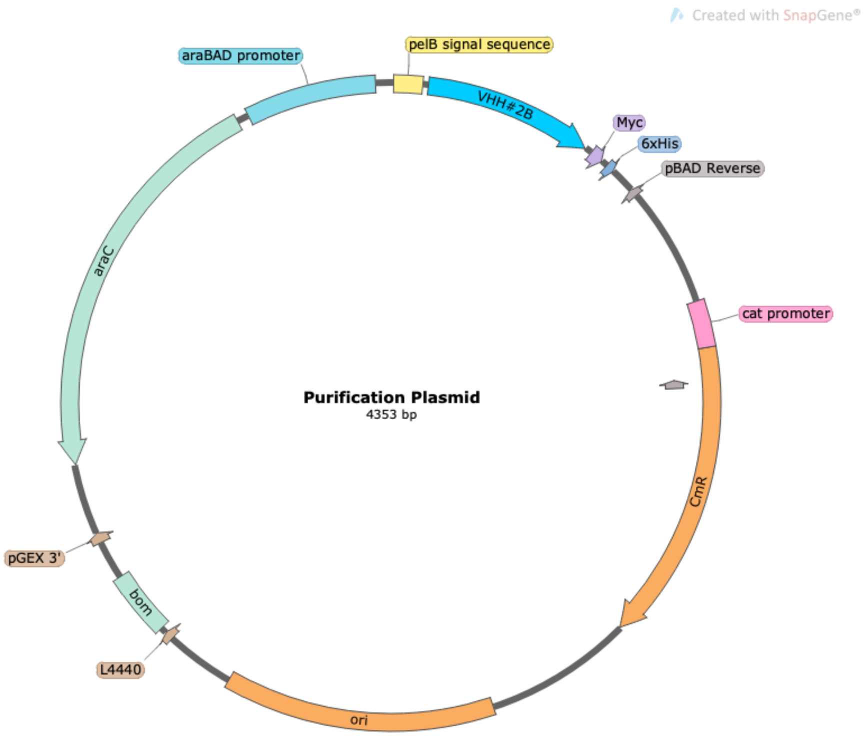


**5.6. Supplementary figure S6. Plasmid map of the nanobody purification plasmid.** The purification plasmid encodes a high copy number origin (colE1), a FX cloning site allowing the exchange of the protein of interest to be purified, a Myc- and His-tag which is automatically added to the protein upon successful integration, and the inducible araBAD promoter with the corresponding araC gene. Additionally, a pelB signal is incorporated, allowing the directed protein transportation to the bacterial periplasm.

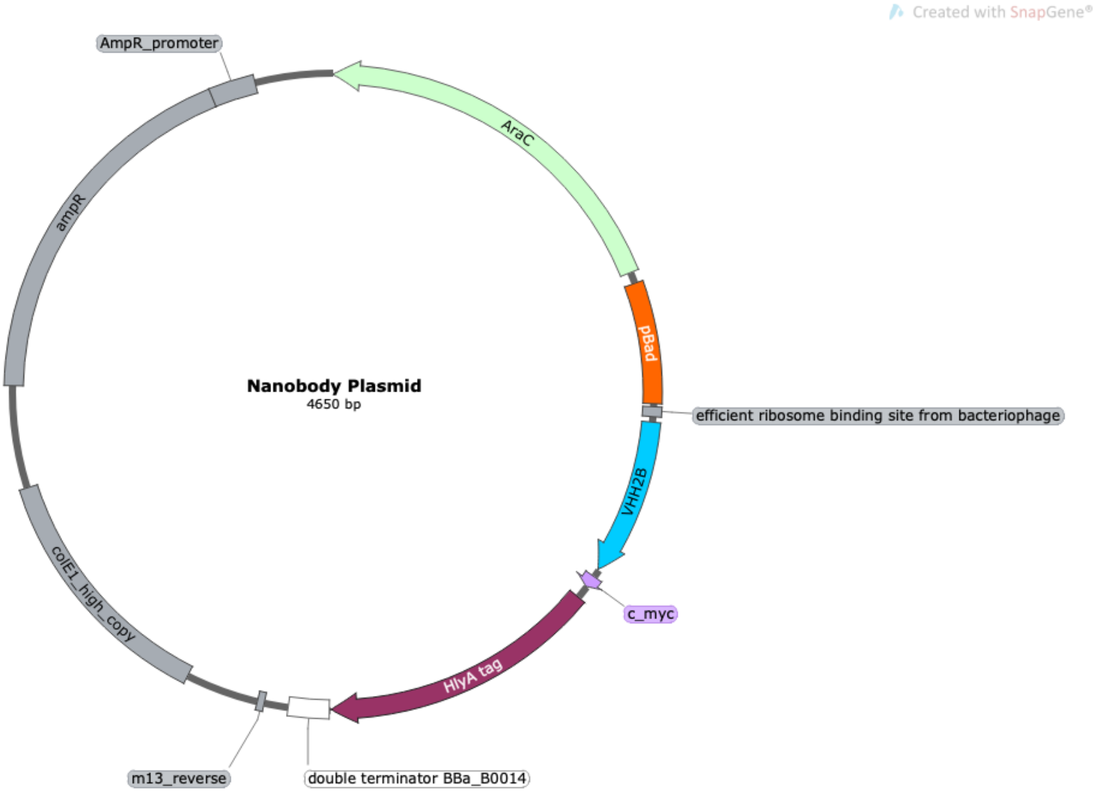


**5.7. Supplementary figure S7. Plasmid map of arabinose-induced nanobody expression plasmid.** The nanobody plasmid encodes a high copy number origin (colE1), an interchangeable region flanked by two SapI sites for exchanging the protein of interest, a Myc- and HlyA-tag which is automatically added to the protein upon successful exchange, and the inducible araBAD promoter with the corresponding araC gene.

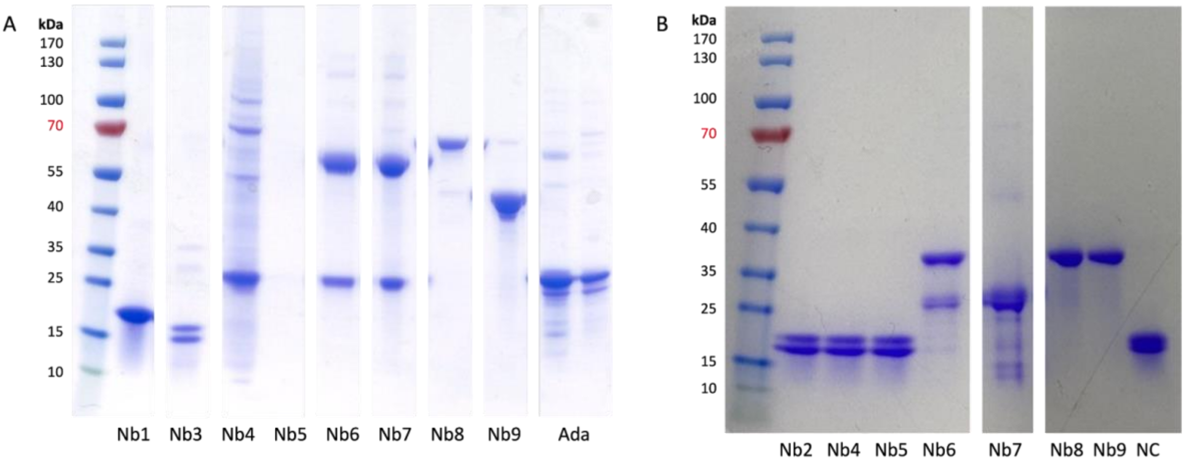


**5.8. Supplementary figure S8. Purification of monovalent and bivalent anti-TNFα nanobodies from *E. coli MC1061* a. Periplasmic extraction performed for all nanobodies.** Periplasmic extraction was being particularly impactful on bivalent nanobody constructs, where the harsh conditions of the extraction led to the breakage of the linkers between coupled nanobodies. Additionally, the figure showcases the production of Adalimumab (Ada) in HEK213 cells, followed by its purification using immobilized metal anion chromatography (IMAC). **b. Periplasmic extraction only performed for monovalent nanobodies and whole cell lysis for bivalent ones.** Whole cell lysis is more suitable to purify bivalent nanobodies.

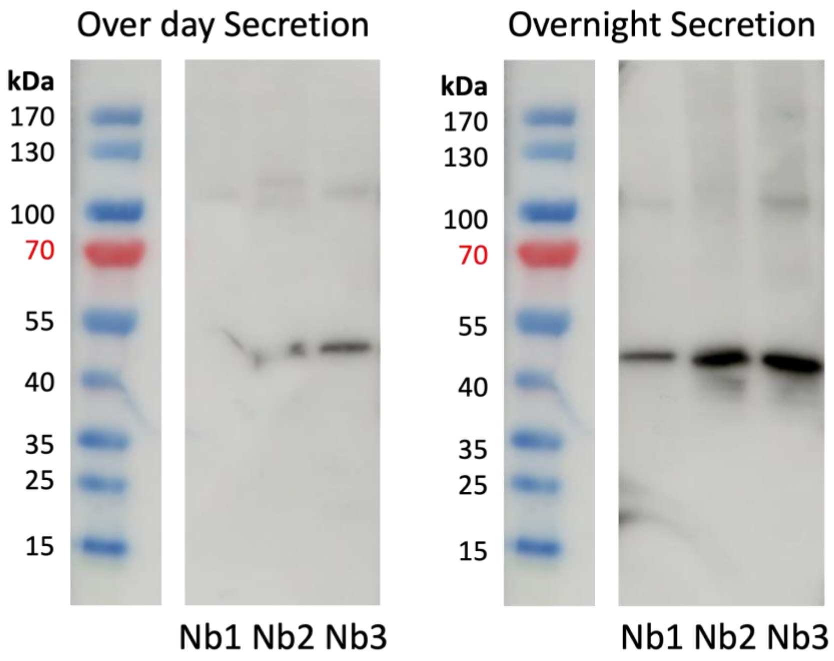


**5.9. Supplementary figure S9. Comparison of over day to overnight arabinose-induced nanobody secretion in *E. coli MC1061*.** Double transformed *E. coli MC1061* were induced by arabinose and incubated at 37°C either over day for 5 hours or overnight for approximately 15 hours. In order to receive enough nanobodies for further testing the overnight induced nanobody expression was continued to be used for the following experiments.

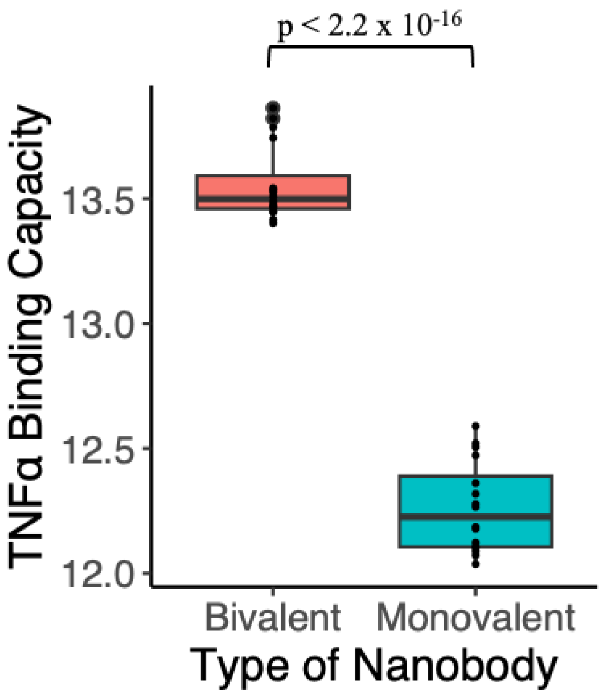


**5.10. Supplementary figure S10. Comparison of TNFα binding capacity between monovalent and bivalent nanobodies.** The boxplots illustrate the binding capacity of TNFα to monovalent and bivalent nanobodies. The red boxplot represents the set of bivalent nanobodies which demonstrate a mean binding capacity of 13.5 ± 0.1, while the teal boxplot represents the set of monovalent nanobodies with a mean binding capacity of 12.3 ± 0.2. A Welch Two Sample t-test reveals a highly significant difference in binding capacities between the groups (t = 21.915, df = 29, p-value < 2.2e-16), with a 95% confidence interval for the difference in means ranging from 1.175 to 1.416. These results robustly support the superiority in TNFα binding of bivalent constructs over their monovalent counterparts.

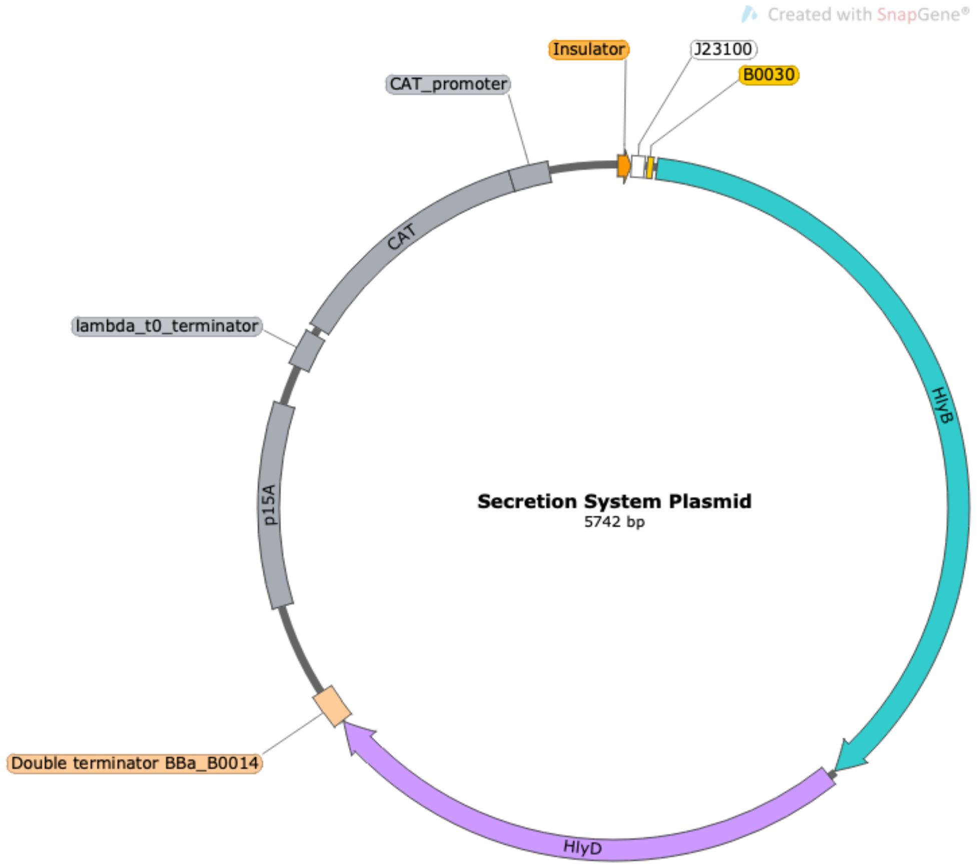


**5.11. Supplementary figure S11. Plasmid map of the secretion system plasmid**. The secretion system plasmid encodes a low copy number origin (p15A), the HlyB and HlyD genes required for the functionality of the one-step secretion system, the constitutive J23100 promoter, and an interchangeable region flanked by two BsmbI-v2 sites for exchanging the promoter in order to regulate further the expression of the secretion system machinery.

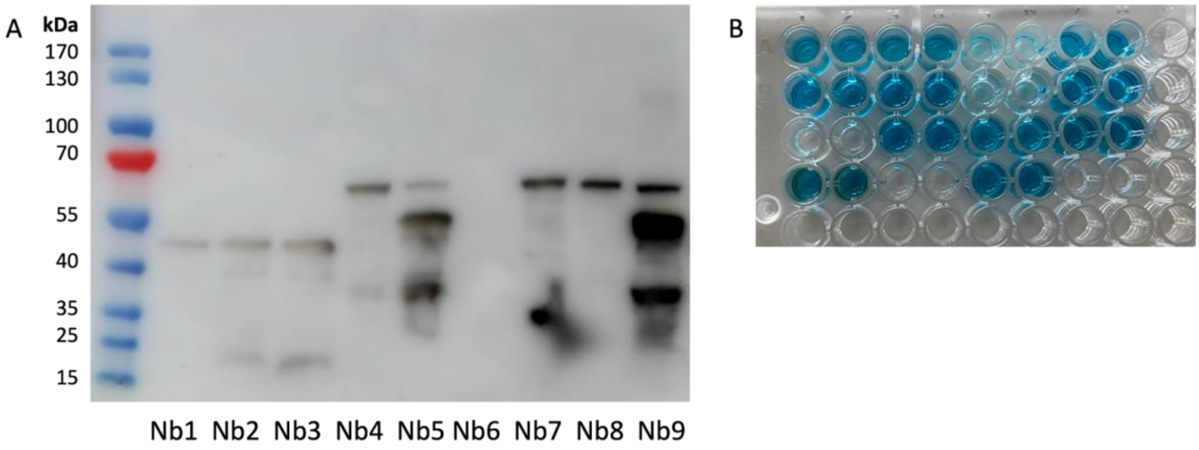


**5.12. Supplementary figure S12. Arabinose-induced anti-TNFα nanobody secretion in *E. coli MC1061***. **a.** Double transformed *E. coli MC1061* were induced by arabinose and incubated at 37°C overnight. Anti-myc antibodies were used in the Western blot to detect secreted nanobodies in the bacterial supernatant. **b. ELISA comparing the TNFα-binding capabilities of secreted vs purified nanobodies obtained from *E. coli MC1061*.**

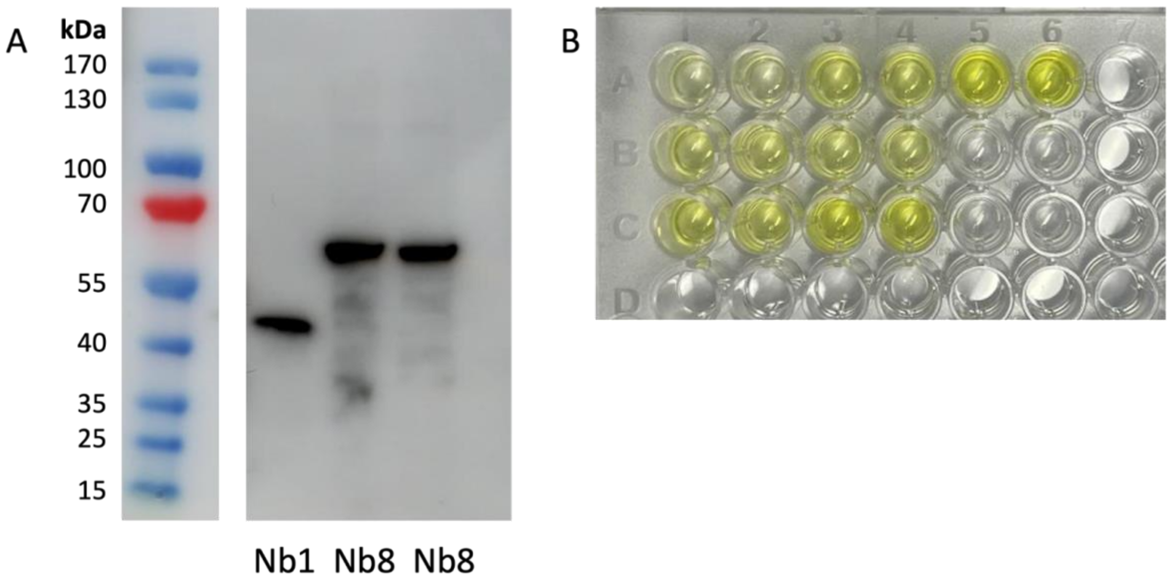


**5.13. Supplementary figure S13. Arabinose-induced anti-TNFα nanobody secretion in *E. coli Nissle 1917*. a.** Double transformed EcN were induced by arabinose and incubated at 37°C overnight. Anti-myc antibodies were used in the Western blot to detect secreted nanobodies in the bacterial supernatant. **b. ELISA displaying the TNFα-binding capabilities of secreted nanobodies obtained from EcN.**

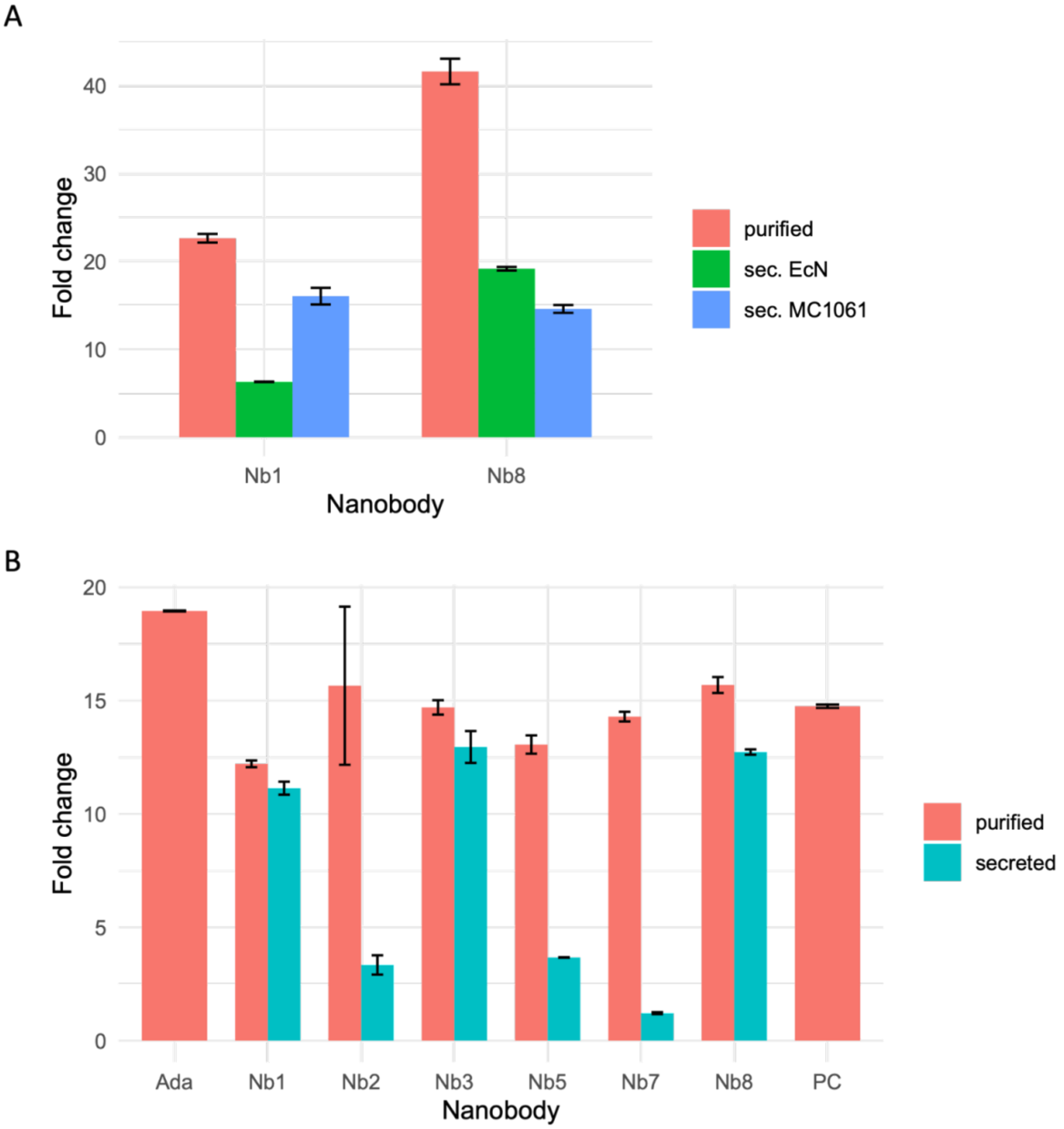


**5.14. Supplementary figure S14. Analysis of ELISA comparing the binding capabilities of purified and secreted monovalent and bivalent anti-TNFα nanobodies in *E. coli Nissle 1917* and *E. coli MC1061* a.** Comparison of secreted anti-TNFα nanobodies in MC1061 and EcN to purified nanobodies obtained from MC1061. **B.** Comparison of the binding capability of purified and secreted anti-TNFα nanobodies obtained from *E. coli MC1061*.

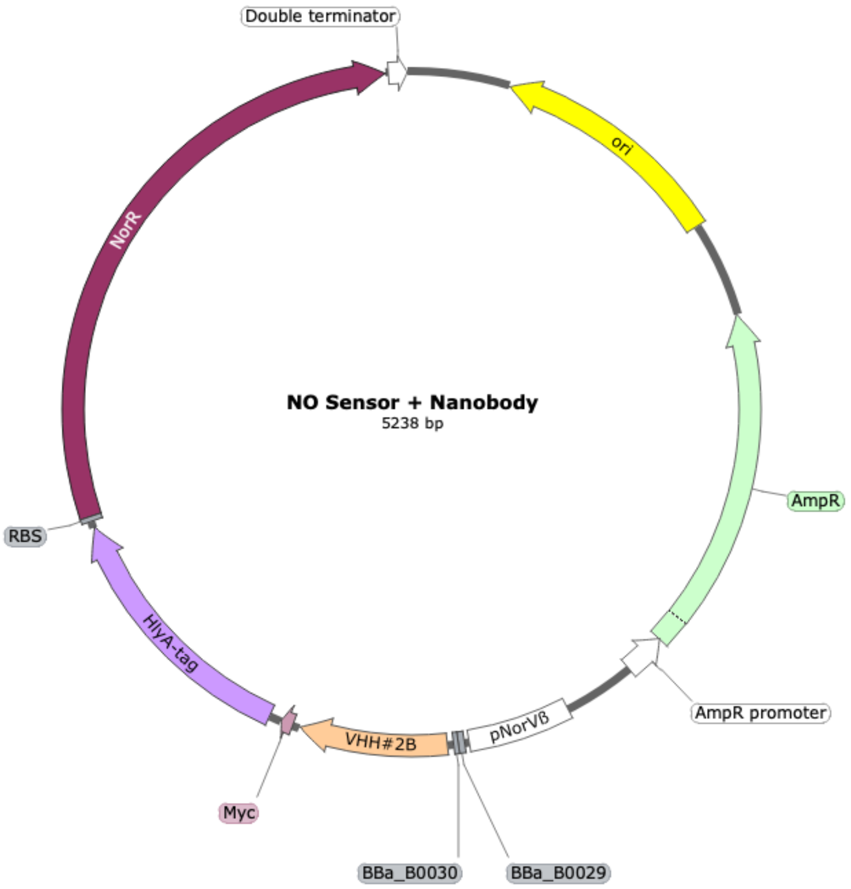


**5.15. Supplementary figure S15. Plasmid map of NO-induced nanobody expression system (ß-2)**. The NO sensor + nanobody plasmid encodes a high copy number origin (colE1), the monovalent nanobody Nb1 with a Myc- and HlyA-tag, and the inducible pNorVß NO sensor with its corresponding NorR gene for the positive feedback loop. This plasmid map displays the ß-2 construct containing 2 RBS in front of the nanobody.

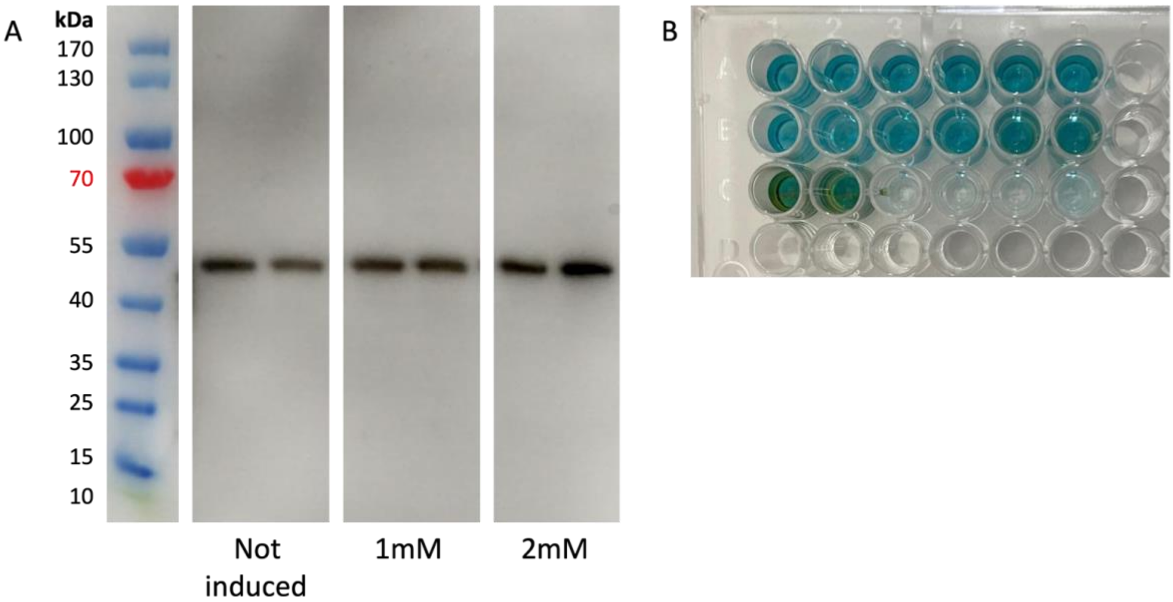


**5.16. Supplementary figure S16. NO-induced monovalent anti-TNFα nanobody secretion and function in *E. coli Nissle 1917.* a.** Double transformed EcN were induced with NO and incubated at 37°C overnight. The expression of the nanobody was under the control of a two-RBS system (ß-2) which showed stronger responses and higher production but also high expression leakage. **b.** ELISA showing the TNFα-binding capabilities of the secreted monovalent nanobody VHH#2B (Nb1) upon NO induction, obtained from EcN.

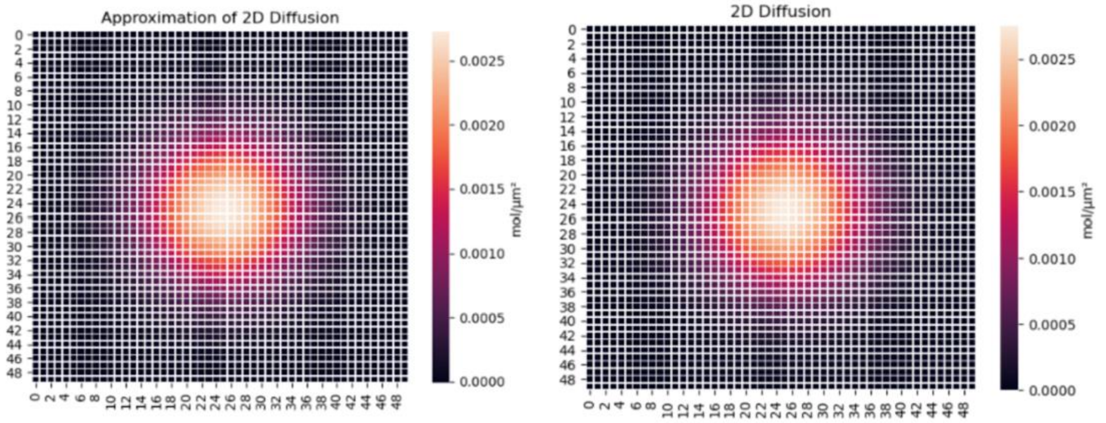


**5.17 Supplementary figure S17. Visual comparison between diffusion models**

Diffusion model used in the final model (left) next to diffusion by the more accurate Crank-Nicolson scheme paired with the Runge-Kutta scheme.
